# PARP1 Inhibition Halts EBV+ Lymphoma Progression by Disrupting the EBNA2/MYC Axis

**DOI:** 10.1101/2023.07.05.547847

**Authors:** Giorgia Napoletani, Samantha S. Soldan, Toshitha Kannan, Sarah Preston-Alp, Peter Vogel, Davide Maestri, Lisa Beatrice Caruso, Andrew Kossenkov, Asher Sobotka, Paul M. Lieberman, Italo Tempera

**Affiliations:** The Wistar Institute, Philadelphia, PA, 19104, USA; Department of Comparative Pathology, St. Jude Children’s Research Hospital, Memphis, TN, USA

**Keywords:** EBV-associated cancer, lymphoma, PARP1, BMN-673, MYC

## Abstract

PARP1 has been shown to regulate EBV latency. However, the therapeutic effect of PARP1 inhibitors on EBV+ lymphomagenesis has not yet been explored. Here, we show that PARPi BMN-673 has a potent anti-tumor effect on EBV-driven LCL in a mouse xenograft model. We found that PARP1 inhibition induces a dramatic transcriptional reprogramming of LCLs driven largely by the reduction of the *MYC* oncogene expression and dysregulation of MYC targets, both *in vivo and in vitro*. PARP1 inhibition also reduced the expression of viral oncoprotein EBNA2, which we previously demonstrated depends on PARP1 for activation of MYC. Further, we show that PARP1 inhibition blocks the chromatin association of MYC, EBNA2, and tumor suppressor p53. Overall, our study strengthens the central role of PARP1 in EBV malignant transformation and identifies the EBNA2/MYC pathway as a target of PARP1 inhibitors and its utility for the treatment of EBNA2-driven EBV-associated cancers.

**Significance Statement:** A promising approach to treating EBV-driven malignancies involves targeting cancer and EBV biology. However, investigating host factors that co-regulate EBV latent gene expression, such as PARP1, has been incomplete. Our study demonstrates that the PARP1 inhibitor BMN-673 effectively reduces EBV-driven tumors and metastasis in an LCL xenograft model. Additionally, we have identified potential dysregulated mechanisms associated with PARP1 inhibition. These findings strengthen the role of PARP1 in EBV+ lymphomas and establish a link between PARP1 and the EBNA2/MYC axis. This has important implications for developing therapeutic approaches to various EBV-associated malignancies.

## Introduction

The gammaherpesvirus Epstein–Barr virus (EBV) is a common human pathogen, with an estimated prevalence of over 90% of the population worldwide (1). EBV was the first human tumor virus discovered and has been causally associated with several malignancies, including non-Hodgkin Lymphoma (e.g., Burkitt lymphoma), Hodgkin Lymphoma, and Diffuse Large B-cell Lymphoma (DLBCL) (2–4). In B cell lymphomas, the expression of EBV proteins acts as the driving force behind oncogenesis, fueling the progression of the disease. In certain instances, expression of this viral protein contributes to a more aggressive tumor behavior in EBV-associated malignancies when compared to non-EBV tumors. Despite the significant role of viral infection in tumor development, we currently lack a specific therapy that directly targets EBV-driven oncogenesis. As a result, EBV-associated malignancies are treated using the same approaches as those not associated with EBV, which highlights the need for more tailored treatments.

In both EBV-associated malignancies and EBV+ cancer cell lines, the Epstein-Barr virus (EBV) expresses a specific set of viral genes without generating viral particles. These viral genes, known as latent viral genes, are expressed by EBV in various gene expression programs referred to as latency types. These latency types are specific to certain EBV+ malignancies or particular stages of infected B-cell differentiation, showcasing distinct patterns of gene expression.

Overall, the fully immortalized B-cells express five EBNAs, two LMPs, EBV-encoded small RNAs (e.g. EBERs), and non-coding BART (Bam HI-A region rightward transcripts) RNAs (5–8). The full expression of viral latent genes is commonly found in B-cell lymphomas that develop in immunosuppressed patients, and it is also characteristic of immortalized lymphoblastoid cell lines (LCLs) *in vitro*. The EBV latent genes code for proteins that play a crucial role in the establishment and maintenance of a persistent infection.

EBNA2 is the main transcription factor of EBV and is expressed at the early stage upon infection, orchestrating the B cells transformation by prompting changes in cell metabolism and stimulating cell proliferation pathways (6, 9–14). EBNA2 has been found to directly activate the transcription of the *MYC* gene in EBV-infected B cells. The dysregulation of MYC expression through EBNA2 contributes to the aberrant cell growth and survival observed in EBV-associated malignancies, particularly B-cell lymphomas. The EBNA2-mediated activation of MYC is considered an important mechanism by which EBV exerts its oncogenic potential. Other viral latent proteins, including LMPs, EBNA3s, and EBNA-LP. can be expressed in tumors and support EBNA2’s transactivating role in cellular survival and proliferation. However, in immunocompetent individuals, EBV eludes immune system detection by adopting a very restrictive latency program characterized by the exclusive expression of noncoding RNAs and EBNA1 (8). *In vivo*, however, sporadic low-level viral reactivations might occur during the host lifespan, causing the induction of lytic genes expression such as *BZLF1* which encodes the lytic transactivator factor Zta, and *BMRF1*, encoding for the polymerase associated factor EA-D (15–19). Indeed, the ability of the EBV to regulate its own gene expression allows it to establish latent infections in host cells and is a crucial factor in the development of EBV-associated malignancies. Targeting the mechanisms that govern EBV latent gene expression holds great potential as a therapeutic strategy for specific treatment of EBV-associated malignancies. By identifying drugs or interventions that specifically interfere with the viral gene expression machinery, it may be possible to disrupt the survival and growth of EBV-associated malignancies.

EBV viral expression is strictly regulated by several host factors, including the poly(ADP-ribose) polymerase 1 (PARP1). PARP1 transfers poly(ADP-ribose) (PAR) moieties (PARylation) on itself and its targets, causing conformational alterations that also result in functional changes (20). PARP1 is a multifaceted host enzyme, playing a central role in transcription regulation, DNA repair, and cell metabolism (20–22). In the last decades, considering its role in DNA damage response, PARP1 has arisen as a critical therapeutic target in several types of cancer, especially those harboring mutations in other DNA repair pathways (22–27). Our group has previously identified how PARP1 can control EBV latency by: (a) altering the 3D virus chromatin structure (28); (b) regulating CTCF binding on EBV promoters and supporting the latency expression program (29–32); (c) repressing the lytic gene expression by binding *BZLF1* promoter (33, 34). To date, the therapeutic effect of PARP1 inhibitors on EBV+ lymphomagenesis has been poorly explored. Therefore, we aimed to investigate whether PARP1 inhibition (PARP1i) would be able to counteract EBV-driven tumors in a LCL xenograft model, and identify and confirm possible mechanisms underlying its therapeutic effect. In the present study we demonstrate that PARP1i restricts EBV-driven lymphoma *in vivo*, pointing out the oncogene *MYC* as its functional target. Specifically, PARP1 inhibition reduces tumor growth and the metastatic potential of EBV+ LCL, inducing a dramatic transcriptional reprogramming. Interestingly, the absence of PARP1 activity causes a decrease in MYC expression, subsequently leading to a dysregulation of MYC-associated co-factors and targets, both *in vivo* and *in vitro*. Our findings also corroborate the link between PARP1 and EBNA2 expression, that we previously demonstrated *in vitro*. Overall, our study strengthens the central role of PARP1 in EBV malignant transformation and outlines the EBNA2/MYC pathway as an additional target of PARP1 regulation in LCL.

## Results

### PARP1 Inhibition Prevents EBV-driven Tumor Growth and Metastasis in Mice

To test the hypothesis that PARP1i can counteract EBV tumors *in vivo*, we engrafted 16 NSG mice (8 females and 8 males; T_e_) with a lymphoblastoid cell line expressing eLuciferase to monitor their growth using bioluminescence (**Fig. 1A**). After seven days (T_0_), we normalized the cohort by the average flux values of the tumor (measured as photons/second, [p/s]). We assigned n=8 mice per experimental group (4 females and 4 males) to be treated with PARP1i or vehicle (Veh) daily by oral administration. BMN-673 (also known as Talazoparib) was selected as PARP1i given its high inhibitory potential at lower doses (0.33 mg/kg (w/w)) compared to other commercially available compounds (35). For the duration of treatment, animals were monitored daily. We observed no significant effect on weight or overall health (*SI Appendix,* **Fig. S1A and S1B**) in mice treated with BMN-673 or Veh, except for subject loss in the Veh group at T_27_ (*SI Appendix*, **Fig. S1C**). At T_28_, mice were sacrificed, and tumors and livers were collected. We observed a significant increase in Total flux [p/s] in the Veh group compared to the PARP1 inhibitor-treated group (**Fig. 1B and 1C**). To evaluate the tumor burden, we also considered the tumor growth inhibition (TGI%) induced by BMN-673 with respect to the vehicle treatment. Results showed that PARP1 inhibition induced a significant reduction of the tumor burden (TGI%= 80.85% ± 4.15 (SEM)) compared to the control group, corroborated by a significant decrease in average radiance (*SI Appendix***, Fig. S1D** and **S1E**). Within the last two weeks of the study, we observed an unexpected spreading of the tumor beyond the area of implantation (the primary tumor), especially in the Veh group (**Figure 1B**); therefore, on the day of the sacrifice, we collected both the primary tumor and the liver, which was the organ consistently positive for eLuciferase signal in the control group (data not shown) and processed the samples for further analysis. Histopathological evaluation of hematoxylin and eosin (H&E) liver staining revealed extensive metastasis in all Veh samples, frequently centered on periportal areas and associated with ischemic necrosis. In contrast, the presence, extent, and severity of neoplastic cell infiltrates were markedly reduced in the BMN-673 mice (**Fig. 1F**). These results were corroborated by immunohistochemical (IHC) staining for the human nuclear mitotic apparatus protein 1 (NUMA1), which specifically labels human cells and permitted quantification of human LCL-driven metastasis in mouse livers. Image analysis of whole slide sections of mouse liver showed a significantly higher percentage of NUMA1-positive cells and NUMA1-positive cell per unit area in the control group than in BMN-673 treated mice (**Fig. 1G**). These findings demonstrated that PARP1i caused a dramatic tumor growth inhibition and a remarkable reduction in the severity and extent of neoplastic infiltrates *in vivo*.

**Figure 1.**
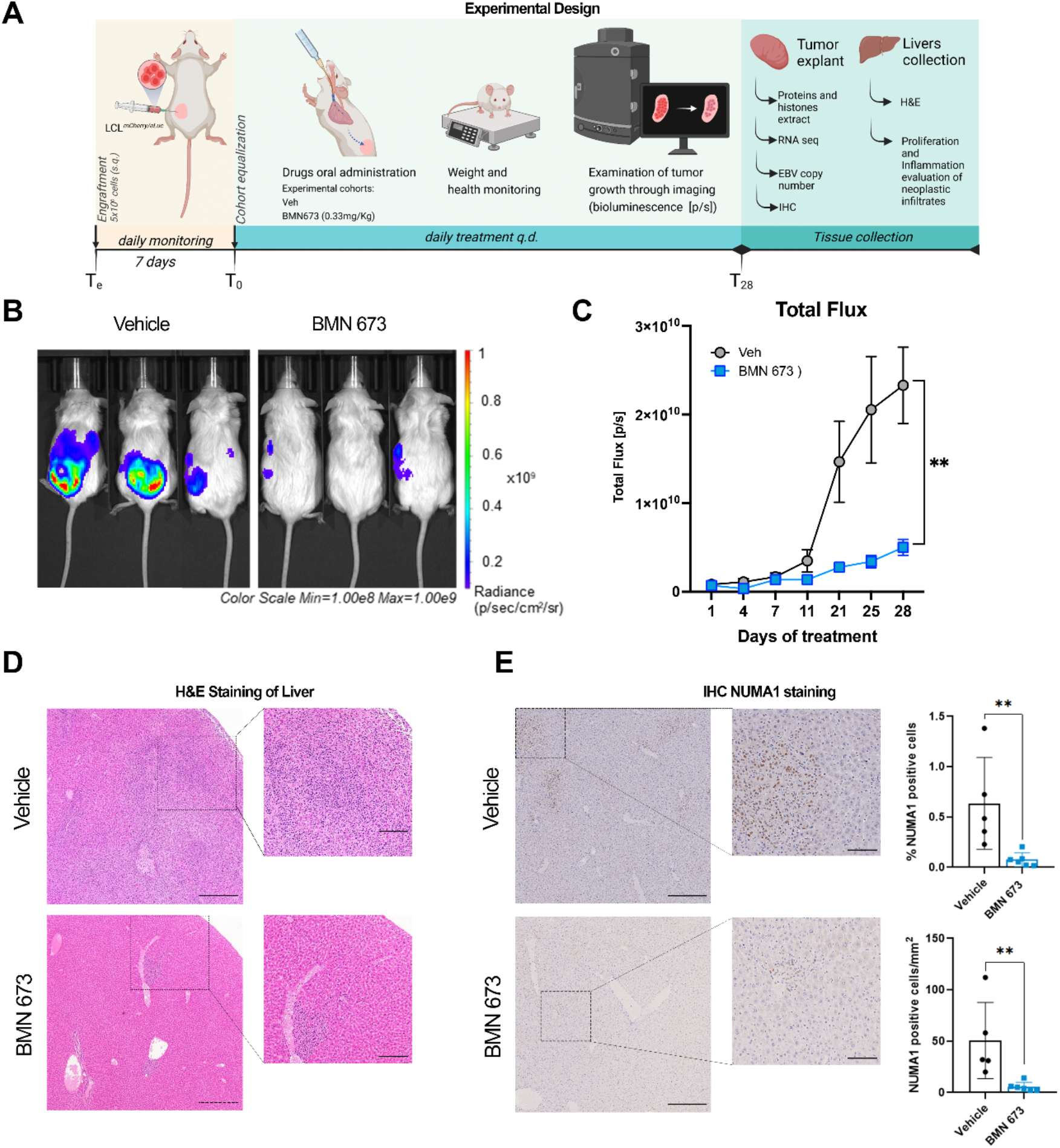
PARP1 inhibition decelerates tumor growth in mice. (**A**) Study experimental design. NSG mice were engrafted with 5x10^6^ LCL expressing eLuciferase (T_e_). After 7 days (T_0_), mice were treated with Vehicle (Veh) or BMN-673 q.d., and tumor growth was monitored by bioluminescence every 2-3 days. On the day of sacrifice (T_28_), the tumors and the livers were explanted for further analyses. (**B**) Representative images of Vehicle and BMN-673 treated mice at T_28_. Bioluminescent signal on each mouse is indicative of the tumor area and intensity. Radiance values are reported as photons/s/cm^2^/sr in a scale from 1x10^8^ (blue) to 1x10^9^ (red). (**C**) Results show the averages ± standard error (SEM) of the Total flux [p/s] values between Veh (gray dots) and BMN-673 -treated (blue dots) mice at T_28_ (n=7 Veh group; n=8 BMN-673 group), analyzed by mixed-effects analysis and Sidak *post-hoc* test (*\*\*p*=0.0012). (**D**) Representative images of livers H&E staining; insets represent the magnification of the LCL metastatic infiltrates visible in disorganized dark purple areas. Nuclei are stained with hematoxylin (dark purple), cytoplasm are stained with eosin (purple). (**E**) Representative images of liver IHC for nuclear NUMA1; positive cells are stained in brown (scale bar: 400µM). Insets represent 4x magnification of metastatic infiltrates (scale bar: 100µM). Results are shown as % of NUMA1 positive cells and NUMA1 positive cells per mm^2^ (n=5 Veh group; n=6 BMN-673 group). Statistical significance has been determined by the Mann-Whitney test, *\*\*p*=0.004).

### PARP1 Inhibition Reduces Global Poly(ADP)ribosylation without Inducing Additional DNA Damage

We previously demonstrated that PARP1 enzymatic activity is essential for EBV gene expression in latency Type III cells, regulating the Cp and BZLF1 promoters, and stabilizing CTCF binding and the chromatin looping across the viral genome (28–31, 33, 34). To assess whether BMN-673 treatment was effective on PARP1 activity in our mouse model, immunofluorescence (IF) staining of PAR and PARP1 was performed. Data showed a significant decrease in nuclear PARylation levels in BMN-673 group compared to Veh, whereas no significant change was found in PARP1 expression (**Fig. 2A**). Additionally, we analyzed the PAR levels in tumor protein extracts by ELISA assay, confirming that PARylation was 3-fold significantly reduced (**Fig. 2B**) in tumors from BMN-673 treated mice. PARP1 plays a central in role in DNA repair (22, 36–38) and PARP inhibitors have been used to elicit DNA damage accumulation in tumors with impaired DNA repair machinery. Therefore, we assessed whether BMN-673 treatment could have caused an increase in DNA damage in tumor samples by measuring via IF the level of phosphorylate H2A.X (γH2A.X), a histone H2A variant that serves as a docking site for DNA damage response and repair factors and therefore used as marker of DNA breaks. Interestingly, we observed γH2A.X positive staining in EBV tumors within the Veh group, indicating that a basal level of DNA damage already exists within these tumors (**Fig. 2C**). In BMN-673 mice, even though we observed a more heterogenous signal in γH2A.X among samples, none of these differences were statistically significant when compared to Veh group (**Fig. 2C**). To further validate IF results, we decided to assess γH2A.X levels in the two groups by western blotting analysis of tumor protein extracts. Consistent with IF analysis, in the Veh group we observed a protein band for γH2A.X in all the tumor samples, supporting the conclusion that basal levels of DNA damage exist in EBV+ malignancies (**Fig. 2D**). In the BMN-673 treated group, we observed again a great variability for γH2A.X signal among the samples, and no significant difference between the two groups was evident with respect to γH2A.X levels (**Fig. 2D**). To confirm that PARP inhibition elicits no accumulation of DNA damage in EBV+ B cells, we determined the amount of DNA damage by *in vitro* single-cell gel electrophoresis (SCGE) in the same cell line implanted before and after treatment with increasing doses of BMN-673 for 72 hours. To control that further DNA damage can be induced in EBV+ LCLs we also assessed DNA damage in cells treated with 20µM of the DNA damaging agent etoposide (ET). Consistent with IF and WB analysis, no significant difference between control (DMSO) and BMN-673 treated cells was observed (**Fig. 2E**); however, a significant increase in DNA damage levels was detected in the samples treated with etoposide (**Fig. 2E**). Overall, these results indicate that PARP inhibition elicits no accumulation of DNA damage in tumor samples from BNM673 treated mice, suggesting that DNA damage was not determinant in causing the observed tumor growth inhibition.

**Figure 2.**
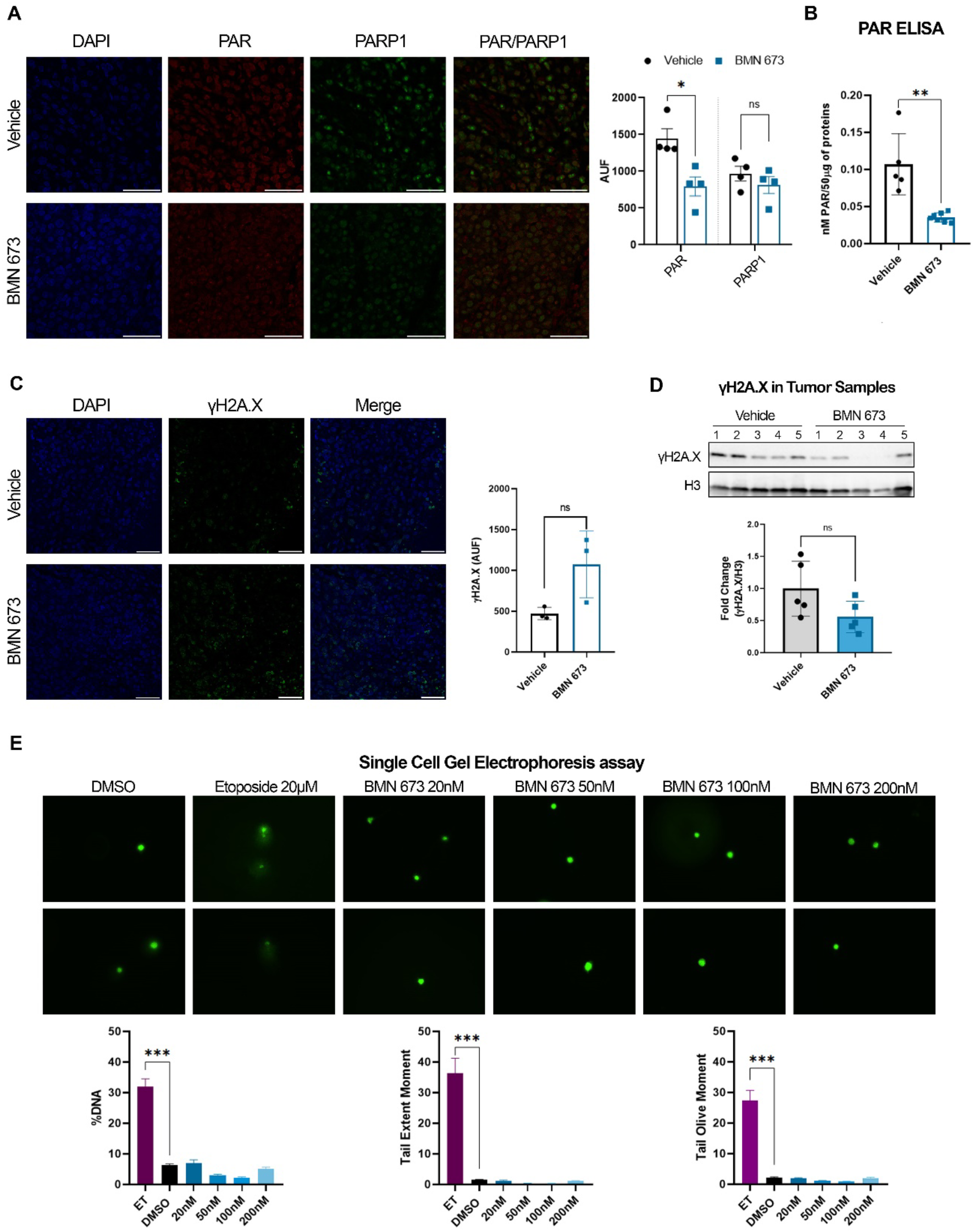
BMN-673 treatment decreases PARylation without increasing DNA damage. (**A**) Representative images of PAR (red) and PARP1 (green) IF staining on tumor sections of Veh and BMN-673 -treated mice (Magnification 63x; scale bar 50µM). Nuclei are stained with DAPI (blue). Results confirmed the decrease of nuclear PARylation and are reported in the bar graph as Raw Intensity (AUF) values (n=4 per experimental group). Statistical significance has been determined by Mann-Whitney test (*p=0.028). (**B**) PAR quantification in tumor total protein extracts by ELISA. Data are showed as nM of PAR for 50µg of protein lysates analyzed in triplicate. Statistical significance has been determined by Mann-Whitney test, **p=0.002. (**C**) Representative IF staining of γH2A.X foci (green) and nuclei (DAPI, blue) on tumor sections (Magnification 40x; scale bar 50µM). Results are shown in the bar plot as average Raw Intensity values ± SEM (n=3 per experimental group). (**D**) Western blot analysis on tumor histones enriched lysates for γH2A.X, confirming the phosphorylation of H2A.X variant in both Veh and BMN-673 tumors (n=5 per experimental group). Histone H3 was used as loading control and data are represented as fold change with respect to Veh values. (**E**) Representative microscopy images of DNA damage analysis by SCGE. LCLs were treated with BMN-673 at a dose of 20 nM, 50nM, 100nM or 200nM for 3 days. Etoposide 20µM was used as positive control to induce DNA damage. DMSO was used as negative control. Bar plots represent the fluorescence intensity average ± SEM of %DNA, Tail extent moment and Tail olive moment. Statistical significance has been determined by One-way ANOVA and Dunnett’s multiple comparison post-hoc test (ET vs DMSO, ***p<0.0001).

### PARP Inhibition Causes Tumor Transcriptional Reprogramming

Beside its role in DNA repair, PARP1 is an essential regulator of gene expression (20, 22, 39) and PARP1-mediated gene regulation is involved in several cellular processes, including EBV-driven gene expression as shown by our group (28, 31, 34, 40). To assess whether and how PARP1 inhibition impinges EBV+ tumor growth observed in our *in vivo* model, we performed RNA-seq on a subset of tumors (Veh n=3; BMN-673 n=4) to evaluate changes in gene expression between control and BMN-673 treated groups. The Principal Component Analysis (PCA) identified that ∼30% of the observed variation in gene expression between the groups is caused by PARP inhibition (**Fig. 3A**), sorting out the experimental group in two different clusters along the Principal Component 1 (PC1) axes. Interestingly, the PC2 axes separated the samples by biological sex, which was more evident in the Veh group compared to BMN-673 treated group, suggesting that the treatment efficacy was unbiased. RNA-seq Transcriptional profiles of Veh and BMN-673 tumors were compared and identified a significant dysregulation of 3112 genes (q<0.05) after PARP inhibition (**Fig. 3C**). Selected hits from the RNA-seq were validated by qPCR (*SI Appendix***, Fig. S2B**). Further analysis showed that BMN-673 differentially expressed genes (DEG) were skewed toward upregulation, with 1807 DEG (58%) showing an increase in expression in the BMN-673 treated group (**Fig. 3C**). Among the top upregulated genes (FC > 2) we observed several histone cluster 1 and 2 transcripts codifying for H2A and H2B (e.g., *HIST1H2AC, HIST1H2BC*, *HIST1H2BJ*, *HIST2H2BE*) isoforms, the transcriptional regulator Early Growth Response 1 (*EGR1*), MAX Dimerization Protein 1 (*MXD1*), and the p53 and p73 suppressors E3 ubiquitin ligase *MDM2* (**Fig. 3C** and *SI Appendix,* **Fig. S2A**). Among the top downregulated genes (FC < -2) after PARP inhibition, we found several genes involved in hematological malignancies including *DOK2,* a scaffold protein associated with chronic myelogenous leukemia, the tumor protein *TP73,* known to have dual and opposite roles in the induction of apoptosis (41). Interestingly, the two oncogenes with a well-established role in lymphomagenesis and EBV+-driven oncogenesis, *MYC* and *MYCL* (11, 42–44) were also among the most downregulated genes after PARP inhibition.

**Figure 3.**
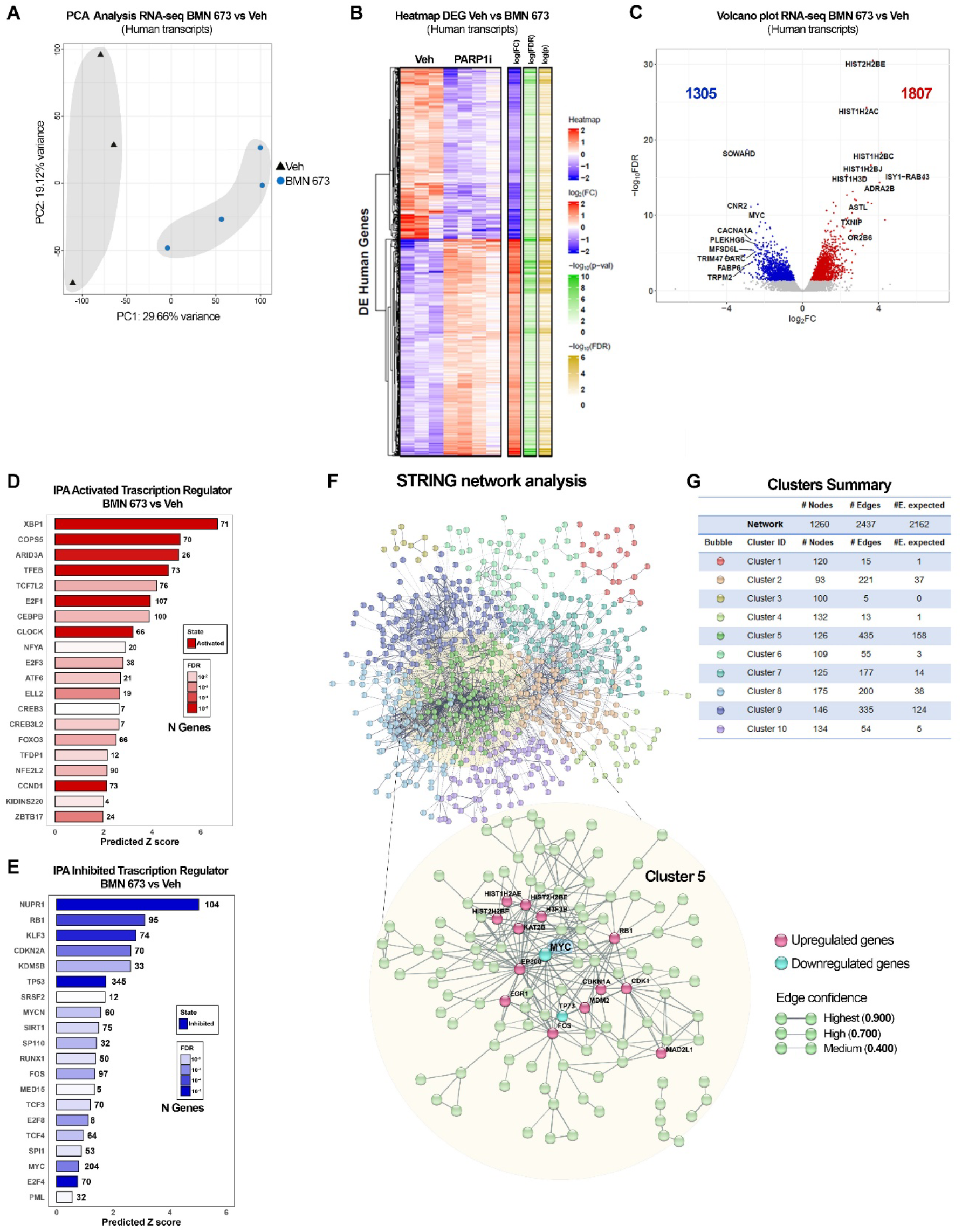
PARP1 inhibition alters human gene expression. (**A**) Principal Component Analysis (PCA) of RNA-seq on Veh and BMN-673 mice (n=3 and n=4, respectively). Samples are separated as a function of Principal Component 1 (PC1, treatment) and PC2 (biological sex). The percentage of variance is indicated on the axes. (**B**) Heatmap of dysregulated human genes expression (DEG) in RNA-seq dataset, after BMN-673 treatment compared to Veh. (C) Volcano plot of the 3112 DEG. The left side of the graft reports the downregulated genes (1305, blue dots) and the right side reports the upregulated genes (1807, red dots). Genes with 2-fold change and false discovery rate FDR < 5% were considered as significantly differentially expressed. The top genes codifying for proteins have been labeled on the plot. (**D**-**E**) Top twenty IPA activated and inhibited transcription regulators, respectively. (F) Representation of DEG STRING network analysis. The ten clusters obtained by k-mean analysis (confidence=0.400) are represented by different colors, as indicated in the network and clusters summary panel (**G**). Magnification shows Cluster 5 interconnections (confidence=0.700), and highlights the downregulated genes in cyan (i.e., MYC, TP73) and upregulated genes in magenta (e.g., EGR1, CDK1).

PARP1 activity affects gene expression also by regulating the functions of several transcription factors. We utilized IPA upstream regulators analysis to determine the activation or the inhibition of transcriptional regulators that could account for the observed changes in gene expression between Veh and BMN-673 groups (**Fig. 3D**). IPA analysis predicted the activation of several transcription factors including XBP1, TFEB, and ARID3A that play a role in B cell differentiation and autophagy and have been linked to EBV infection (**Fig. 3D**). Among the upstream regulators predicted to be inhibited by PARP inhibitors we observed NUPR1, RB1, and TP53 (**Fig. 3E**). Notably, consistent with our RNA-seq analysis, IPA highlighted MYC and MYCN (another member of the MYC family), in the top 20 Inhibited transcriptional regulators, suggesting the hypothesis that differences induced by the PARP inhibitor may be related to MYC downregulation. Overall, these data indicate that PARP inhibition transcriptionally reprogrammed tumor cells and downregulated important oncogenes.

### PARP Inhibition Disrupts MYC-driven Gene Expression Program

MYC has been proven to be pivotal in EBV latency maintenance and EBV-driven lymphomas (11, 43, 45–47), acting as a ubiquitous amplifier of gene transcription and proliferative signaling. To further validate the inhibitory effect of BMN-673 treatment on MYC transcriptional functions we analyzed our RNA-seq data set for the expression of a subgroup of genes regulated by MYC, using as references the two curated human gene sets for MYC gene from the Gene Set Enrichment Analysis (GSEA) (MYC_v1 and MYC_v2). We found that several genes in both MYC signatures were significantly (p<0.05) deregulated by PARP1 inhibition (*SI Appendix,* **Fig. S2C**). Furthermore, to identify networks within the transcriptome, we performed a STRING analysis of the 1260 most DEG genes (FDR<5%, FC|Z|≥2) in our RNA-seq dataset. We filtered the network by functional and physical protein-protein association, co-expression, and co-occurrence in databases, removed all the disconnected nodes from the analysis, and generated a map of all interactions existing between the genes deregulated in tumor samples after PARP inhibition (**Fig. 3F**). To evaluate the nodes in the network and gain a better insight into the biological functions that DEG genes affected, we performed a K-means clustering analysis of the network followed by Gene Ontology (GO) enrichment analysis of biological processes (**Fig. 3G**). We identified ten main clusters differentially enriched in the number of nodes and biological functions. The top hits included “B cell activation involved in immune response”, “Positive regulation of lymphocyte differentiation”, as well as “Intrinsic apoptotic signaling pathway in response to DNA damage by p53 class mediator” (GO:0002312, GO:0045621, GO:0042771, respectively) (*SI Appendix***, Table S1**). Next, we focused our attention on Cluster 5, where most connections occurred (edges=435, nodes=126, expected edges=158) (**Fig. 3G**). A stricter GO analysis on cluster 5 highlighted “transcription” in the top 20 biological functions (**Table 1**). GO analysis showed that the main enriched proteins in terms of the number of biological processes involved within the cluster were represented by E1A binding protein P300 EP300 (235), Cyclin-dependent kinase 1 CDK1 (221), the transcription factor MYC scored (183) and Histone acetyltransferase KAT2B (154) (**Table 2**). Within cluster 5, by adjusting it for a minimum interaction score of confidence=0.7 (high confidence), we retrieved several deregulated genes including MYC (**Fig. 3E**). Given the downregulation of *MYC* transcripts and the inactivation shown by IPA, we used IHC to assess in tumor samples whether the levels of MYC protein were reduced in BMN-673 group compared to control. Consistent with the RNA-seq data, we observed that both the percentage of cells positive for MYC and the number of cells per mm^2^ positive for MYC were significantly decreased in the BMN-673 mice in comparison to Veh (**Fig. 4A**). In lymphomas, the activation of MYC plays a significant role in the spontaneous inactivation of the ARF-Mdm2-p53 pathway (48), therefore to further consolidate the observed transcriptional changes between the two groups, we evaluated the level of p53 after PARP1 inhibition. By IHC staining, we found that expression p53 protein was markedly increased in PARP1i-treated mice compared to control group (**Fig. 4B**). Altogether, these data strongly showed the central role played by MYC in mediating tumor growth inhibition via PARP1 inhibition.

**Figure 4.**
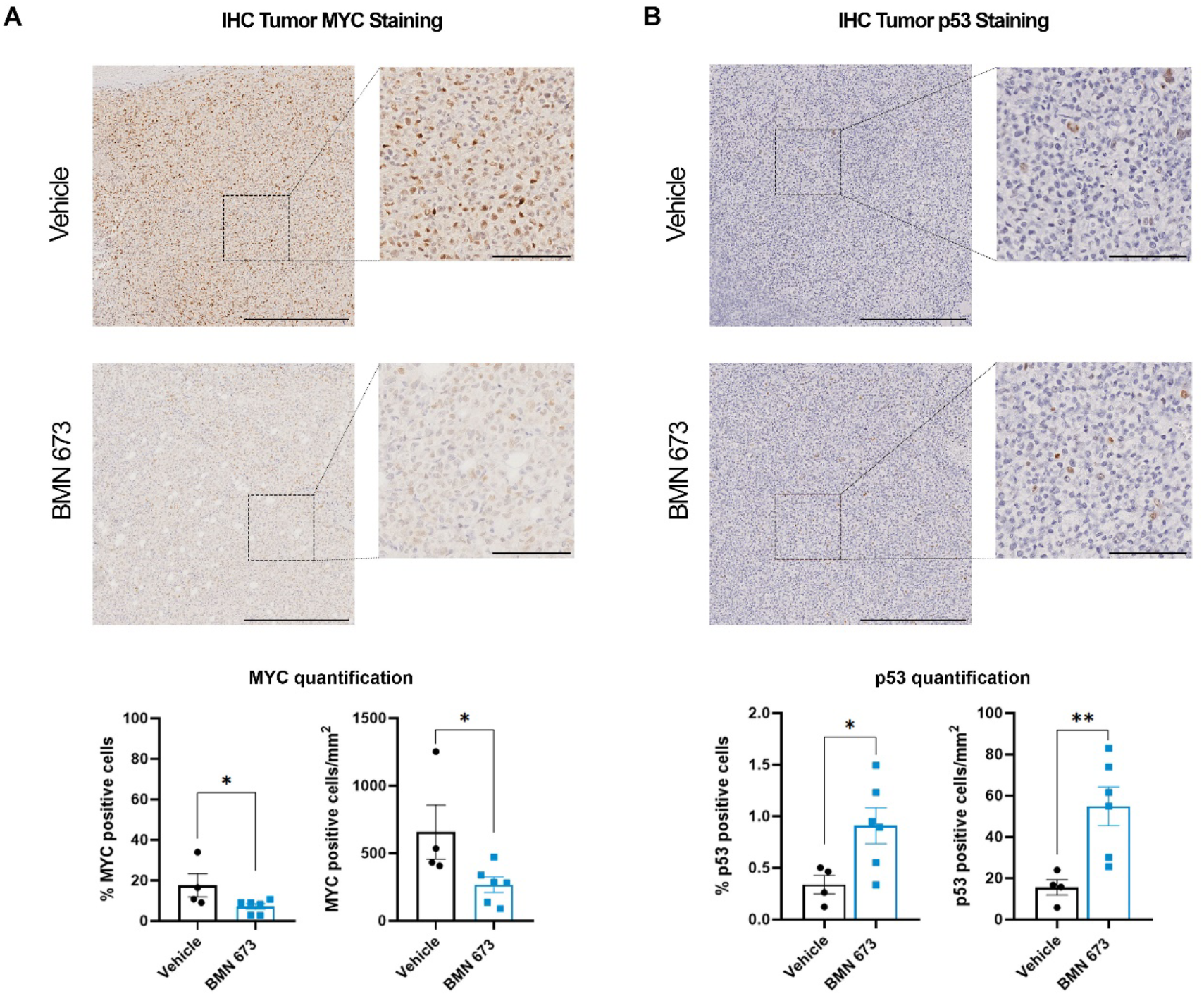
PARP1 inhibition induces changes in MYC and p53 expression in tumor sections. (**A**) Representative IHC staining of MYC (brown) and nuclei (hematoxylin, purple) in tumor sections. Insets are a representative zoomed area of the tumor (4x magnification). (**B**) Representative IHC images of p53 (brown) and nuclei (hematoxylin, purple) in tumor sections. Insets are a representative zoomed area of the tumor (4x magnification). MYC and p53 quantifications were performed by percentage of positive cells (left bar plot) and number of positive cells per area (mm^2^) (n=4 Veh group; n=6 BMN-673 group). Images scale bar 400 µM; insets scale bar 100 µM. Statistical significance has been determined by Mann-Whitney test (%MYC *\*p*=0.010; MYC positive cells/mm^2^ *\*p*=0.038; %p53 *\*p*=0.038; Myc positive cells/mm^2^ *\*\*p*=0.009).

**Table 1.** Cluster 5 GO Biological Functions of STRING Network Analysis Top 20 GO Biological Functions organized by Strenght. In Bold are highlighted MYC-enriched functions

| GO #term ID | GO Biological Functions description | observed<br>gene count | background<br>gene count | Strength | FDR |
| --- | --- | --- | --- | --- | --- |
| GO:0035257 | Nuclear hormone receptor binding | 10 | 155 | 1 | 1.21E-05 |
| **GO:0140297** | **DNA-binding transcription factor binding** | **22** | **366** | **0.97** | **2.84E-12** |
| GO:0061629 | RNA polymerase II-specific DNA-binding transcription factor binding | 15 | 283 | 0.92 | 1.30E-07 |
| **GO:0008134** | **Transcription factor binding** | **33** | **672** | **0.88** | **3.02E-16** |
| GO:0003682 | Chromatin binding | 26 | 570 | 0.85 | 3.05E-12 |
| GO:0031625 | Ubiquitin protein ligase binding | 13 | 296 | 0.83 | 1.14E-05 |
| GO:0003713 | Transcription coactivator activity | 13 | 316 | 0.81 | 2.04E-05 |
| **GO:0001228** | **DNA-binding transcription activator activity, RNA polymerase II-specific** | **16** | **449** | **0.74** | **6.17E-06** |
| GO:0003712 | Transcription coregulator activity | 20 | 571 | 0.74 | 1.69E-07 |
| GO:0016887 | ATPase activity | 14 | 393 | 0.74 | 3.54E-05 |
| **GO:0000977** | **RNA polymerase II transcription regulatory region sequence-specific DNA binding** | **30** | **878** | **0.72** | **1.67E-11** |
| GO:0046982 | Protein heterodimerization activity | 11 | 338 | 0.7 | 0.0013 |
| **GO:1990837** | **Sequence-specific double-stranded DNA binding** | **34** | **1068** | **0.69** | **3.05E-12** |
| GO:0000987 | Cis-regulatory region sequence-specific DNA binding | 22 | 701 | 0.69 | 1.69E-07 |
| **GO:0000978** | **RNA polymerase II cis-regulatory region sequence-specific DNA binding** | **21** | **672** | **0.69** | **3.98E-07** |
| GO:0003690 | Double-stranded DNA binding | 36 | 1156 | 0.68 | 1.19E-12 |
| GO:0000976 | Transcription regulatory region sequence-specific DNA binding | 32 | 1028 | 0.68 | 2.28E-11 |
| GO:0043565 | Sequence-specific DNA binding | 39 | 1331 | 0.66 | 6.72E-13 |
| GO:0000981 | DNA-binding transcription factor activity, RNA polymerase II-specific | 30 | 1022 | 0.66 | 5.55E-10 |
| GO:0140110 | Transcription regulator activity | 47 | 1657 | 0.64 | 1.13E-15 |

**Table 2.** Cluster 5 GO Biological Processes of STRING Network Analysis Top 20 GO Biological Processes organized by Strenght. In Bold are highlighted MYC-enriched Processes

| GO #term ID | GO Biological Processes description | observed gene count | background gene count | Strength | FDR |
| --- | --- | --- | --- | --- | --- |
| GO:0006367 | Transcription initiation from RNA polymerase II promoter | 20 | 162 | 1.28 | 9.85E-17 |
| GO:0006352 | DNA-templated transcription, initiation | 22 | 214 | 1.2 | 5.31E-17 |
| GO:0006366 | Transcription by RNA polymerase II | 26 | 406 | 1 | 1.03E-15 |
| **GO:0045637** | **Regulation of myeloid cell differentiation** | **15** | **260** | **0.95** | **3.26E-08** |
| GO:0097659 | Nucleic acid-templated transcription | 30 | 568 | 0.91 | 3.14E-16 |
| GO:0006351 | Transcription, DNA-templated | 29 | 567 | 0.9 | 2.68E-15 |
| GO:0048545 | Response to steroid hormone | 15 | 328 | 0.85 | 5.35E-07 |
| **GO:1903706** | **Regulation of hemopoiesis** | **20** | **493** | **0.8** | **1.29E-08** |
| GO:0032870 | Cellular response to hormone stimulus | 22 | 569 | 0.78 | 3.79E-09 |
| GO:0045787 | Positive regulation of cell cycle | 15 | 388 | 0.78 | 4.12E-06 |
| **GO:0000122** | **Negative regulation of transcription by RNA polymerase II** | **34** | **895** | **0.77** | **1.38E-14** |
| GO:0061061 | Muscle structure development | 18 | 479 | 0.77 | 2.96E-07 |
| **GO:0045786** | **Negative regulation of cell cycle** | **21** | **571** | **0.76** | **2.26E-08** |
| GO:0071407 | Cellular response to organic cyclic compound | 20 | 537 | 0.76 | 4.65E-08 |
| **GO:0045944** | **Positive regulation of transcription by RNA polymerase II** | **44** | **1253** | **0.74** | **2.01E-18** |
| GO:0009725 | Response to hormone | 30 | 849 | 0.74 | 6.60E-12 |
| GO:0048608 | Reproductive structure development | 15 | 420 | 0.74 | 1.00E-05 |
| **GO:0000278** | **Mitotic cell cycle** | **24** | **695** | **0.73** | **4.01E-09** |
| **GO:0006325** | **Chromatin organization** | **24** | **713** | **0.72** | **6.29E-09** |
| **GO:1903047** | **Mitotic cell cycle process** | **21** | **616** | **0.72** | **7.40E-08** |

### PARPi Treatment *in vitro* Results in Reduced MYC Levels, Mirroring Findings in a Mouse Model of EBV-positive Tumors

Our experiments *in vivo* indicate that PARP inhibition interferes with MYC functions. To gain a better insight into the link between PARP inhibition and MYC at the mechanistic level and its implication in EBV-driven lymphomagenesis, we decide to further assess the effect of PARP inhibition *in vitro*, using the same LCL cells we implanted in mice. First, we evaluated the IC50 of BMN-673 using CellTiterGlow assay (**Fig. 5A**). We treated LCLs with different concentrations of BMN-673 for either 3 or 5 days; we selected a longer time point to determine the effects of PARP inhibition after multiple rounds of replication. We established the BMN-673 EC50 of ∼200 or 300 nM for 3 days and 5 days of treatment, respectively (**Fig. 5A and** *SI Appendix,* **S3A)**. To further characterize BMN-673 cytotoxicity and to determine which mechanism of cell death was activated by this PARP inhibitor, we assessed AnnexinV and propidium iodide (PI) (**Fig. 5B**). FACS analysis showed that BMN-673 globally induced cell death (Q_1-3_) in ∼67% of cells at the EC50 dose of 200nM after 72hrs, and this effect was exacerbated in treating cells up to 5 days (*SI Appendix,* **Fig. S3B**). Specifically, the number of cells positive for both AnnexinV^+^/PI^+^, which indicated cells in late-stage apoptosis or already dead, was ∼35% (Q_2_), and for the AnnexinV staining (Q_3_), indicating an early-apoptosis event, was ∼30%. Cell death was significantly reduced at the lower concentration of 50nM (Q_1-3_ 59%, Q_2_ 28%, Q_3_ 27%) and was reduced to 20% at 20nM, whereas merely necrosis events were observable (Q_1_ 9%). These results indicate that PARP1 inhibition triggers cell death mostly through activation of apoptosis at higher doses of BMN-673, whereas 20nM dose was better tolerated. Next, we investigated at the molecular level the effects of PARP inhibition on LCL cells. *In vivo*, our RNA-seq analysis indicates that PARP1 inhibition interferes with MYC and p53 regulation of gene expression (**Fig. 3D**). MYC and p53 regulate gene expression by binding to the regulatory regions of target genes, which requires them to associate with chromatin. PARP1 activity regulates the interactions between transcription factors and chromatin, therefore we hypothesized that PARP inhibition may affect MYC and p53 interactions with chromatin. Moreover, the apoptosis events that we previously observed might be downstream of p53 activation. To assess the effects of PARP inhibition on the chromatin localization of MYC, p53, and PARP1, we analyzed nuclear soluble (SN) and chromatin-bound (CB) fractions of LCL cells before and 72 hours after treatment with increasing concentrations of BMN-673. Our results showed that inhibiting PARP increased the levels of MYC, p53, and PARP1 in the nuclear soluble fraction (**Fig. 5C**). We only detected MYC, p53, and PARP1 proteins in the Chromatin Bound (CB) fraction in the untreated (DMSO) samples. There was no significant signal detected for these proteins in the CB fractions after PARP1 inhibition (**Fig. 5C**). In addition, we also evaluated PARylation levels in the SN and CB fractions before and after PARP1 inhibition using Dot-Blot analysis. Interestingly, our results showed that treatment with BMN-673 completely abolished PARylation in the CB fraction (**Fig. 5D**). However, in the SN fraction, PARylation levels were significantly decreased in a dose-dependent manner with BMN-673 treatment, but some PARylation was still detected (**Fig. 5D**). These findings indicate that PARP1 inhibition reduces the association of MYC and p53 with chromatin, impairing their ability to regulate gene expression. Additionally, the data showed that in LCL cells BMN-673 treatment did not induce PARP1 trapping unless used at high concentrations, providing evidence for a DNA-repair independent role of PARP1 in tumor growth. These observations add further interest to our study. *In vivo*, we determined that PARP1 inhibition also affected MYC and p53 protein levels. To confirm these findings in an *in vitro* setting, we conducted Western blot analysis to evaluate MYC and p53 levels in LCL cells before and after treatment with BMN-673. Our results showed that PARP inhibition significantly reduced MYC levels while increasing p53 levels (**Fig. 5E**), which is consistent with our *in vivo* observations.

**Figure 5.**
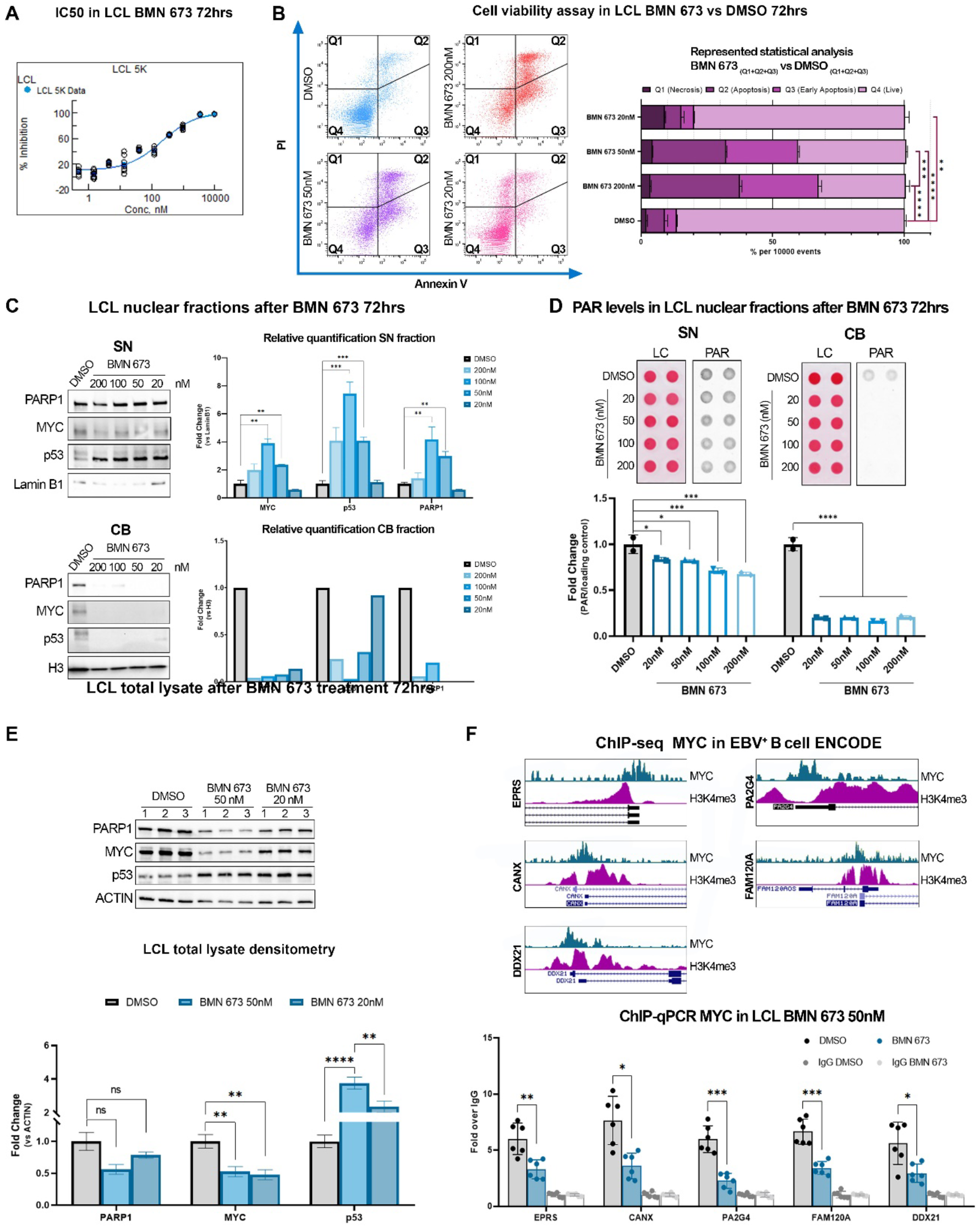
BMN-673 treatment causes MYC depletion in LCL. (**A**) BMN-673 IC50 after 3 days of treatment. Resultant EC50 is ∼200 nM. (**B**) AnnexinV/PI analysis shows the LCL triplicates treated with DMSO (control) or 20 nM, 50nM, or 200nM BMN-673 for 3 days. Ten-thousand events were collected per sample. Quadrants detail PI (Q_1_, necrosis), AnnexinV/PI (Q_2_, apoptosis), AnnexinV (Q_3_, early-apoptosis) positive cells, or negative cells (Q_4_, live cells). Statistical analysis was performed on the average ± SEM of BMN-673 compared to the DMSO dead cells quadrants (Q1-3) values by 2-way ANOVA with Tukey *post-hoc* (**p=0.004, ***p<0.001, ****p<0.0001). (**C**) Western blot analysis of PARP1, MYC or p53 in Soluble Nuclear (SN) and Chromatin Bound (CB) fractions. Lamin B1 and histone H3 were used as SN and CB loading controls, respectively. Statistical significance was determined by multiple *t-test* and two-stage step-up FDR method (MYC_100nM_ *\*\*q*=0.008, MYC_50nM_ *\*\*q*=0.005; p53_100nM_ and p53_50nM_ *\*\*\*q*<0.001; PARP1_100nM_ *\*\*q*=0.008, PARP1_50nM_ *\*\*q*=0.001). (**D**) Dot blot analysis of PARylation in nuclear fractions. PAR levels were normalized to the loading control (LC, ponceau). Statistical analysis was performed on the average ± SEM of BMN-673 with respect to DMSO values by 2-way ANOVA and Tukey *post-hoc* (SN, 20nM *\*p*=0.020, 50nM *\*p*=0.013, 100nM and 200nM *\*\*\*p*<0.001, CB, *\*p*<0.0001). (**E**) Western blot analysis of PARP1, MYC and p53 on LCL treated with 50nM or 20nM BMN-673 for 72hrs. Actin was used as loading control. Statistical significance was determined by multiple *t-test* (MYC_50nM_ *\*\*p*=0.009, MYC_20nM_ *\*\*p*=0.005; p53_50nM_ *\*\*\*\*p*<0.0001, p53_20nM_ *\*\*\*p*=0.004). (**F**) ChIP-seq signatures of MYC and H3K9me3 in publicly available datasets on EBV^+^ B cells. Representative tracks show MYC promoter occupancy of those genes dysregulated by PARP1 inhibition in the GSEA signature. (**G**) Quantitative chromatin immunoprecipitation (ChIP-qPCR) evaluation of human DEG represented in (**F**). Results are represented as the average ± SEM fold-change over ChIP-qPCR negative control (IgG). Statistical significance has been determined by one-way ANOVA with Dunnett’s T3 *post-hoc* (EPRS *\*p*=0.008; CANX *\*p*=0.010; PA2G4 *\*\*\*p*<0.001; FAM120A *\*\*\*p*<0.001; DDX21 *\*p*=0.037).

To determine whether the observed changes in MYC translate in a loss of MYC on the promoter of target genes, we assessed MYC occupancy at the promoter of BMN-673 affected genes. For this analysis, we selected a subset of genes that were deregulated by PARP1 in our *in vivo* studies and that were annotated as MYC targets in the curated GSEA dataset (**Fig. S2**). We validated the occupancy of MYC and the active chromatin signature (H3K4me3 deposition) on the promoter of the selected genes using publicly available ChIP-seq datasets for MYC in EBV-infected LCL cells (GSE36354 and GSM945188, respectively) (**Fig. 5F**). We further investigated MYC occupancy at the promoter of these genes in LCL cells before and after BMN-673 treatment using quantitative ChIP analysis. Our results demonstrated that MYC occupancy at the promoter of all tested genes was significantly reduced after treatment with BMN-673 (**Fig. 5G**). Taken together, our results suggest that PARP1 inhibition impairs the ability of MYC to directly associate with chromatin and activate gene expression at the transcriptional level.

### PARP Inhibition Alters the Expression of Epstein-Barr Virus (EBV) Genes

In EBV-positive lymphoma, MYC plays a critical role in the cancer phenotype as its transcription is activated and regulated by the viral protein EBNA2. Our recent findings indicate that PARP1 and its enzymatic activity are essential for the expression of EBV latent genes, including EBNA2. In light of this, we aimed to investigate whether the reduction in tumor growth and dysregulation of human genes we observed *in vivo* correlated with changes in EBV gene expression. To achieve this, we analyzed our RNA-seq dataset to determine the expression of EBV genes. Interestingly, we found that the samples were separated based on BMN-673 treatment by PCA analysis, suggesting that PARP inhibition significantly impacts EBV gene expression (**Fig. 6A**). Next, we determined which viral genes were affected by PARP inhibition. We observed a significant increase in EBV genes generally associated with lytic reactivation, including the lytic genes transactivator *BRLF1* and the polymerase-associated factor *BMRF1,* which code for the early antigen D factor (EA-D) (**Fig. 6B**). Likewise, the oncogene *BARF1* and the virion proteins *BFRF3* and *BFLF1* were also significantly upregulated, together with the nuclear egress factors *BFRF1* and *2.* In contrast, the BART family transcript *A73* was the only gene significantly downregulated after PARP inhibition (**Fig. 6B**). These findings suggest that PARP1 inhibition induces significant changes in the expression of EBV genes. To confirm that PARP1 inhibition represses latent gene expression and promotes expression of lytic genes, we assessed the expression profile of viral genes in the vehicle group. We observed that LMPs and EBNAs genes were the highest viral transcripts, confirming that EBV adopted the latency III program in the vehicle group (**Fig. 6C**).

**Figure 6.**
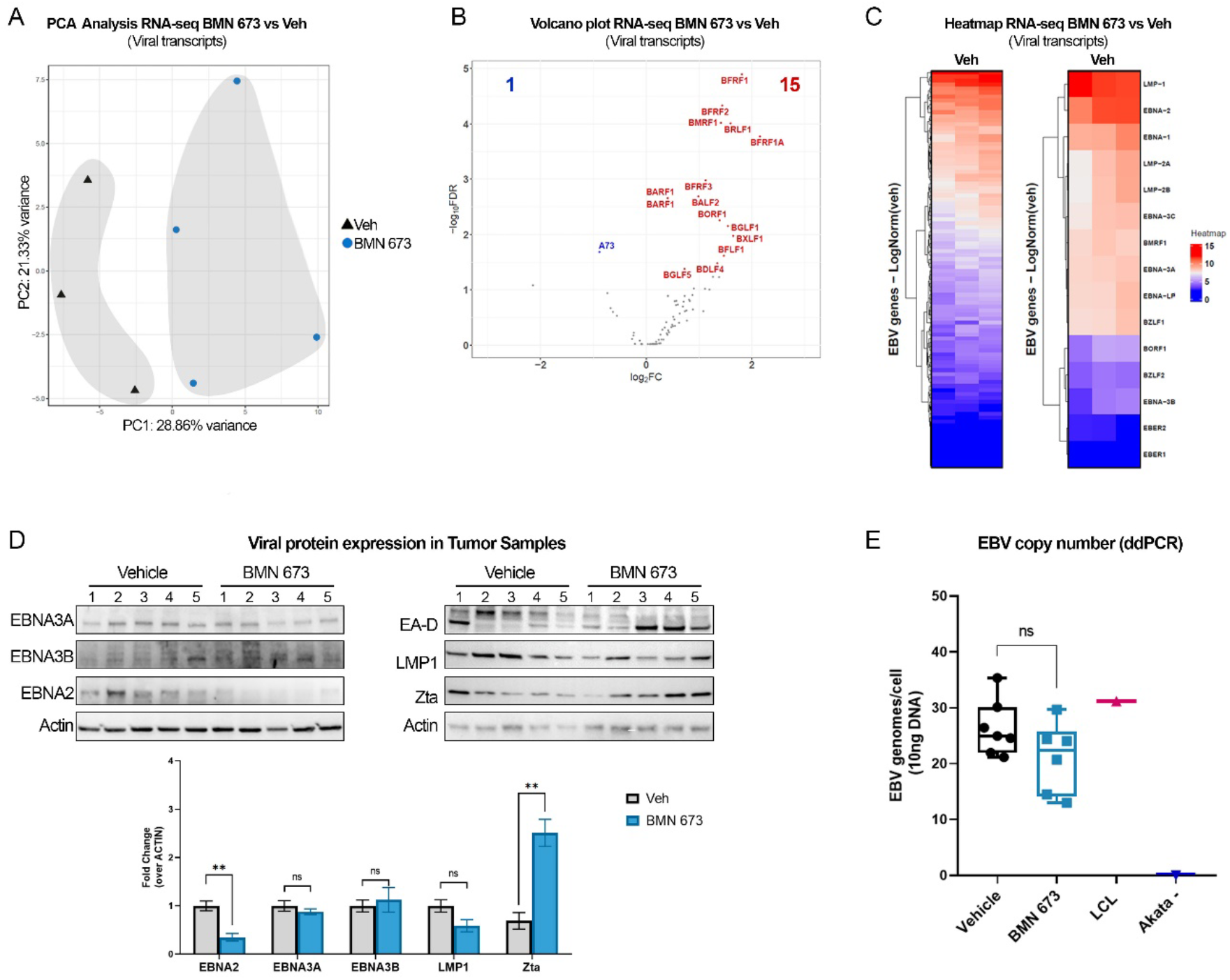
PARP1 inhibition dysregulates EBV gene expression. (**A**) Principal Component Analysis (PCA) of EBV transcripts in the RNA-seq dataset (Veh n=3; BMN-673 n=4). Samples are represented as a function of Principal Component 1 (PC1, treatment) and PC2 (biological sex). The percentage of variance is indicated on the axes. (**B**) Volcano plot of the 16 viral DEG. Genes with 2-fold change and false discovery rate FDR < 5% were considered significantly differentially expressed and labeled on the plot. All the genes that did not pass the FDR<5% cut-off are shown as gray dots. The left side of the graft reports the downregulated genes (A73, blue) and the right side reports the upregulated genes (15, red dots). (**C**) Heatmap of EBV genes expression of Veh group, highlighting EBV latency and lytic-associated transcripts (right panel). (**D**) Western blot analysis of EBV proteins in tumor samples. The expression of the representative latency III (i.e., EBNA2, EBNA3A, EBNA3B, LMP1) and lytic reactivation-associated (i.e., EA-D, Zta) proteins have been tested. Actin was used as loading control. Relative viral protein expression is reported as fold change of BMN-673 over Veh (n=5 per experimental group). Statistical significance was determined by multiple *t-test* (EBNA2, *\*\*p*=0.001; Zta *\*\*p*=0.001). (**E**) EBV copy number quantification. Viral genomes per 10ng of DNA loaded were analyzed in duplicate by digital droplets PCR (ddPCR) in Veh and BMN-673 DNA extracts. LCL EBV^+^ and AKATA EBV^-^ cell lines were used as positive and negative controls, respectively.

Next, we investigated whether the transcriptional changes in viral genes were reflected in changes in viral protein expression in the tumor samples from the Veh and BMN-673 groups. We used western blot analysis to measure the levels of latent proteins (EBNA2, EBNA3A, EBNA3B, and LMP1) and lytic viral proteins (Zta and EA-D) (**Fig. 6D**). We found a significant reduction in the levels of EBNA2 protein in the BMN-673 group compared to the Veh group, consistent with our previous findings of decreased EBNA2 levels after PARP1 inhibition in LCL cells (40) (**Fig. 6D**). Interestingly, no significant changes were observed in the levels of EBNA3A, EBNA3B, or LMP1 proteins after PARP1 inhibition (**Fig. 6D**). We also observed that while Zta protein was detected in both the Veh and BMN-673 groups, its levels were significantly higher in the BMN-673 group compared to the Veh group (**Fig. 6D**). However, we observed variable expression of the lytic protein EA-D in both experimental groups, suggesting that although BMN-673 induces lytic transcripts, these changes are not sufficient to support a productive lytic replication. Complete lytic replication is characterized by a significant increase in the copies of EBV genome per cell. To determine whether BMN-673 treatment induces lytic replication, we accurately measured the number of EBV genome copies per cell in tissue samples from both groups using digital droplet PCR. We observed that infected cells in both groups had a similar number of copies of the EBV genome (**Fig. 6E**), which is consistent with what was observed in LCL cells. Based on the observed changes in EBV gene expression, protein levels, and genome copies per cell, our findings suggest that PARP1 plays a critical role in maintaining EBV latency and its inhibition leads to dysregulation of viral expression that may trigger early, but abortive lytic reactivation. Overall, our study provides insight into the potential use of PARP1 inhibitors in treating EBV-associated malignancies by altering both viral and host gene expression, and ultimately, reprogramming cancer gene expression.

## Discussion

We reported previously that PARP1 plays a critical role in regulating and maintaining EBV latency. While its importance has been established in latent viral infection, the efficacy of PARP1 inhibitors in restricting EBV+ lymphomas and associated lymphoproliferative malignancies remains uncertain. In this study, we aimed to determine the activity of the PARP1 inhibitor BMN-673 on a mouse model of EBV-driven B cell lymphoma. BMN-673 is an oral, highly potent inhibitor approved by the US FDA for the treatment of advanced or metastatic breast cancer (24, 25, 35, 49, 50) and has been shown to interfere with EBV-induced primary B cells transformation in our previous work (40). Our current study found that PARP inhibition successfully reduces EBV-driven lymphoma growth *in vivo*. Remarkably, we report that the treatment with PARP inhibitor significantly reduced neoplastic infiltration in other tissues without any evident impairment of overall mouse health. Mechanistically, we determine that PARP1 inhibition drives transcriptional changes in EBV+ tumors that reduces MYC and MYC-driven gene expression. Our findings suggest that continuous administration of PARP1 inhibitors significantly restricts the growth and propagation of tumors, making it a viable therapeutic option for patients with EBV-driven lymphomas.

PARP1 is a crucial enzyme involved in DNA damage response, playing a pivotal role in repairing single-stranded break (SSBs) and double-stranded break (DSBs) (22). As a cancer therapeutic strategy, PARP inhibitors have been used to induce synthetic lethality by triggering significant DNA damage in tumors that carry mutations in DNA repair proteins. Surprisingly, although we observed that PARP1 inhibition significantly slows the tumor growth in treated mice, we found no significantly appreciable differences in DNA damage levels in EBV+ tumors obtained from the group treated with BMN-673 compared to those from untreated animals. However, we did observe signs of DNA breaks in tumors from the control group, indicating that a source of DNA damage already exists in EBV-positive malignancies. This finding is consistent with previous research showing that EBV infection induces DNA damage. For example, the EBV protein EBNA1 has been shown to cause genomic instability and oxidative stress in infected cells and promote genomic rearrangement (51). Interestingly, we previously reported that PARP1 binds to and modifies EBNA1, which affects its functions (52). Thus, PARP1 may regulate DNA damage response by multiple mechanisms during EBV latency.

PARP inhibitors are known to elicit cytotoxicity in cancer cells by trapping PARP1 on chromatin, but we did not observe an accumulation of PARP1 on chromatin in our *in vitro* analysis of PARP1 levels before and after inhibition. This is somewhat surprising and suggests that in EBV+ lymphomas, PARP inhibitor induced cytotoxicity through mechanisms independent of PARP1 trapping. While we could only assess PARP1 association with chromatin in cultured cells, our data suggest that DNA breaks exist within EBV+ cancer cells but the accumulation of DNA damage is not the primary mechanism through which PARP1 inhibition counteracted tumor growth or EBV+ B cell proliferation. Overall, our findings highlight the complex nature of EBV+ lymphomas and their relationship to DNA damage and PARP1 activity, suggesting that there may be alternative pathways that contribute to the efficacy of PARP inhibitors in treating EBV-driven malignancies.

Recent studies have highlighted the importance of PARP1 and PARylation in regulating gene expression (20). PARP1-mediated gene transcription has been implicated in various cellular processes, including viral infection. For instance, we previously reported that EBV infection activates PARP1 through the viral protein LMP1, and LMP1-induced gene expression requires PARP1 (32). Our transcriptomic data support the notion that PARP1 plays a critical role in controlling gene expression, as we observed a significant change in the transcriptome of tumor cells from the group treated with the PARP inhibitor. Our transcriptomic analysis and IPA analysis revealed the surprising finding that MYC was significantly downregulated in EBV+ tumors treated with PARP inhibitors. MYC is a well-known player in B cell malignant transformation, and it has been demonstrated that the mutual relationship between EBV and *MYC* expression accelerates lymphomagenesis in EBV+ B cells (10, 43, 45, 47, 53). MYC is a transcription factor crucial in promoting cell transformation, and its interaction with chromatin is essential for its function. We observed a significant reduction in MYC protein levels in tumors treated with BMN-673 that correlated with changes in the expression of MYC targets, suggesting that *in vivo* PARP1 inhibition interferes with MYC-regulated gene expression. This hypothesis is supported by our *in vitro* data clearly showing that MYC’s ability to bind to the promoter region of target genes after PARP1 activity is impaired, leading to the deregulation of downstream targets. Our study suggests that targeting PARP1 could be a promising therapeutic approach to counteracting MYC dysregulation in EBV-driven lymphomas. However, our findings are in contrast with previous studies (54), which showed that PARP1 deletion promotes B-cell lymphoma in Eµ-Myc mice, exacerbating tumorigenesis. Notably, the development of lymphoma via PARP1 depletion was only observed in conjunction with Myc overexpression. One potential explanation for this discrepancy is that in our model MYC overexpression is directly linked to viral infection, whereas in Eµ-Myc mice overexpression of Myc is driven by a transgene mimicking the characteristic human Burkitt lymphoma t(8:14) translocation of c*Myc* and *IgH* regulatory elements (55). Thus, in our model, where MYC gene regulatory element is intact (like EBV+ PTLPD), PARP1 inhibition interferes with EBV-mediated regulation of MYC. This hypothesis is consistent with our previous observations showed that PARP1 inhibition represses EBNA2, the viral activator of MYC (28, 30). Therefore, we speculate that PARP1 indirectly controls MYC through the viral oncoprotein EBNA2. Our hypothesis is further supported by previous findings showing that PARP1 inhibitors alter the chromatin landscape of the EBV epigenome and thus reduce EBNA2 expression in two different EBV+ B cell lines (28, 40). Our data *in vivo* confirm that PARP1 is necessary to regulate EBV gene expression epigenetically. Our RNA-seq analysis showed that PARP inhibition significantly changes the viral gene expression program adopted by EBV in the tumor cells, with an increase in the expression of viral genes usually associated with lytic reactivation. This is also in accord with recent observations showing that depletion of MYC promotes EBV lytic reactivation through changes in EBV chromatin structure (43). Changes in viral chromatin structure that induce lytic gene expression were also reported in EBV+ B cells *in vitro* after PARP1 inhibition by our group, further supporting the importance of PARP1 in regulating viral latency (28). However, despite an increasing expression of lytic viral genes, we observed no increase in EBV viral copies or a significant viral reactivation in tumor samples after PARP1 inhibition. The observed expression of lytic genes without full viral replication in our tumor samples is reminiscent of the EBV gene expression observed upon primary infection, where EBV briefly undergoes a pre-latent abortive lytic cycle (19, 56–58), in which some lytic and latent genes are expressed without the production of viral particles. The pre-latent abortive lytic cycle is then resolved by chromatinization of viral episome, establishing EBV latency. Therefore, a possible explanation of our results is that PARP1 activity is critical for the chromatinization of EBV episome and resolution of the pre-latent abortive lytic cycle, and thus, inhibition of PARP1 may promote initiation of the abortive lytic cycle. Given limitations on our xenograft model, further studies are needed to determine whether PARP1 inhibition causes productive or abortive viral reactivation in a model permissive to re-infection, i.e., models with a functional immune system. Our findings suggest that PARP1-mediated gene transcription is a crucial component of EBV-driven lymphoma development and progression by regulating both viral and host gene expression. The essential role of PARP1-mediated gene expression is further supported by our observations that the histone clusters 1 and 2 genes (HIST1, HIST2) are among the most up-regulated transcripts in the BMN-673 -treated groups compared to the control group (**Fig. 3C** and *SI Appendix,* **Fig. S2A**). H1 mutations are highly diffuse in lymphomas arising from germinal center B cells. Melnick and his group recently highlighted Histone H1 isoforms as tumor suppressors in lymphomagenesis (59). Albeit future investigations are needed to characterize how PARP1 controls and regulates histone expression, our findings suggest a potential novel relationship between PARP1 and histones in lymphoma.

Overall, the present work and previous findings reveal a more complex effect of PARP1 inhibitors in EBV-driven B cell transformation. In summary, our data support PARP1 as an effective target *in vivo* for treating EBV+ lymphoma and suggest that this therapeutic effect may be mediated by counteracting the activation of MYC by EBNA2. Our results shed new light on the potential of PARP1 inhibitors as a therapeutic option for EBV-associated lymphomas and highlight the importance of further translational research in this area.

## Materials and Methods

### Cell culture and drug treatment

NHC1 lymphoblastoid cell lines (LCLs) harboring EBV B95.8 strain used in this study were cultured in 15% fetal bovine serum (FBS), 1% Penicillin/Streptomycin (Gibco) RPMI 1640 (Corning) at 37 °C and 5% CO_2_. To assess the half maximal inhibitory concentration (IC50), LCL were treated with PARP1 inhibitor BMN-673 (Talazoparib, LT-673; Selleck Chemicals, Cat. No. S7048) for 72 hours (h), 5 days or 7 days at several doses (serial dilution from 10µM to 0.01 nM), or DMSO (MilliporeSigma, Cat. No. D8418). For the successive experiment, BMN-673 was used at 200nM, 100nM, 50nM and 20nM. PARP1 activity was validated by DB, as detailed below. For SCGE, cell was treated with 20 µM Etoposide (MilliporeSigma, Cat. No. E1383) for 4 h.

### Cell viability, apoptosis and SCGE assays

To determine the BMN-673 IC50, LCL were seeded at different concentrations (10, 5, 2.5 or 1× 10^4^) in 384-well plates. Cell growth inhibition was determined by CellTiterGlow/Resazurin and analyzed on GraphPad. FITC Annexin V PI Apoptosis Detection Kit (BioLegend, Cat. No. 640914) was used to study the apoptosis induction caused by BMN-673 following manufacturer instructions. Briefly, 10 × 10^5^ LCL were treated with DMSO or 200nM, 100nM, 50nM or 20nM BMN-673. After 72 h or 5 days, cells were washed with Cell Staining Buffer and resuspended in Annexin V binding buffer, supplemented with Annexin V/PI and incubated 15 min at RT in the dark. FACS analysis was performed on LSR II-14 flow cytometer (BD Biosciences). Negative control, Annexin V and PI positive staining were used to set the gating, excluding cellular debris. Data were collected and analyzed using FlowJo (BD Biosciences). SCGE was performed on DMSO, BMN-673 and Etoposide treated cells using the Comet Assay Kit protocol (Abcam; Cat. No. ab238544), with minor optimizations. Specifically, LCL were lysate overnight at 4°C, and the electrophoresis was carried out in TBE for 20 min (60, 61). Images were taken at the Nikon TE2000 inverted microscope (20x magnification) using NIS software (Nikon Instruments Inc.) and analyzed by CometAnalyser (62). GraphPad software was used for statistical analysis.

### Mouse xenograft model and treatment

Sixteen 8-weeks-old NSG mice (8 female and 8 male) were subcutaneously injected with 5x10^6^ LCL cells (EBV B95.8 strain) resuspended in cold 20% Matrigel in PBS w/o Ca^2+^/Mg^2+^(Corning). Mice were anesthetized using 2% isoflurane prior to and during the implantation. Seven days post-implant (T_0_), mice were examined by IVIS^®^ Spectrum *in vivo* imaging system (PerkinElmer Inc.), and randomly assigned to treatment group (BMN-673) or control group (Veh) (n=8 per group; 4 female and 4 male each). BMN-673 (0.33 mg/Kg, cat. No. S7048, Selleck Chemicals, Houston, TX, USA) or vehicle (10% DMAc, 6% Kolliphor and 84% PBS) was administered by oral gavage q.b. for 28 days (T_28_) as previous described (35) . Twice a week, the tumor growth was measured by average flux (photons/second, [p/s]) using IVIS bioluminescent imaging; the percentage (%) of tumor growth inhibition, considered as the measure of the tumor burden, was calculated at the end of the study as the ratio between TotalFlux[p/s]_BMN673-T28_-TotalFlux[p/s]_BMN673-T0_ and TotalFlux[p/s]_Veh-T28_-TotalFlux[p/s]_Veh-T0_. Each imaging session was performed 15 minutes after intraperitoneal injecting D-Luciferin infusion (working concentration 15mg/ml, dose 10ml/Kg, MilliporeSigma, Merck KGaA, Darmstadt, Germany), considering the eLuciferase average flux (photons/second, [p/s]) *throughout* 2% isoflurane anesthesia. Mice were euthanized by CO_2_ asphyxiation after 28 days of treatment, and tumors and metastasis-positive tissues were harvested and snap-frozen in dry ice for DNA, RNA, proteins, or fixed in 10% formalin for histological analyses. Engrafted mice were daily monitored for any suffering, distress or behavioral changes, or weight loss by measuring total body weight three times weekly for a total of 5 weeks (35 days) of study. All the procedures performed were previously approved by The Wistar Institute Institutional Animal Care and Use Committee (IACUC) under the Animal Welfare Act regulation (protocol title “Targeting Epstein Barr virus-associated lymphomas”; protocol number 201524-v3).

### Western blot analysis

For whole-cell protein extracts, cells and tissues lysis was performed in radioimmunoprecipitation assay (RIPA) buffer (Millipore, Cat. No.) supplemented with 1x protease inhibitor cocktail (PIC, ThermoFisher Scientific) and 1× PARG inhibitor (PDD, Selleck Chemicals, Catalog No. S8862). Tissues were homogenized with TissueLyser II (Qiagen). Homogenates were incubated for 30 min in a thermomixer (4°C) and protein extracts were collected after centrifugation at 14,000 × g for 10 min at 4°C. For nuclear extract fractions, Subcellular Protein Fractionation Kit from (ThermoFisher Scientific, Catalog No. 78840) was used following the manufacturer’s instructions, supplemented with 1× PIC and 1× PDD. For Histone extraction, tissues were first disaggregated with a Dounce homogenizer in 1X Pre-Lysis Buffer, and then processed following EpiQuik™ Total Histone Extraction Kit protocol (Epigentek). Depending on the assay, protein concentration was measured using a bicinchoninic acid (BCA) protein assay (Pierce) or Bradford assay (Bio-Rad, Cat. No. 5000006). Proteins were prepared in 1× Laemmli buffer (Bio-Rad, Cat. No. 1610747) supplemented with β-mercaptoethanol (Sigma-Aldrich) and resolved by electrophoresis on a 4-20% or 8-16% polyacrylamide gradient gel (Mini-Protean TGX, Bio-Rad). Proteins were transferred to nitrocellulose or PVDF membranes (Immobilon-P membrane, Millipore; Biorad). and were blocked in 5% milk or 2.5% BSA in TBS-T for 1 h at RT. Incubation with the designated primary antibodies, reported in Key Resources Table, was performed at 4°C overnight; HRP-coniugated secondary antibodies anti-rabbit, anti-mouse, anti-sheep (Jackson ImmunoResearch Inc.), or anti-rat (Bio-Rad) were incubated 1h at RT. Chemiluminescence signals were acquired via iBright Imaging System (ThermoFisher Scientific).

### Immunohistochemistry and Immunofluorescence

Formalin-fixed tumor and liver samples were paraffin-embedded in blocks and sliced into 4µm sections, as previously reported (63), with minor modifications. Briefly, tissue sections were deparaffinized using xylene and serial ethanol washes. Heat-induced epitope retrieval was performed with Citrate buffer pH 6.0 in a 98°C steamer for 20 min, followed by 3 washes in diH_2_O. IHC and Hematoxylin and Eosin (H&E) staining were performed by The Wistar Institute Histotechnology core. Hematoxylin was used as nuclear control stain. Evaluation of H&E and NUMA1 staining was performed using HALO® Image Analysis Platform. MYC and p53 staining were acquired through NanozoomerS60, and analyzed using QuPath software. Tissue slides for IF were permeabilized and blocked in 5% BSA TBS 0.3% TritonX for 1 h at RT and then incubated with the chosen primary antibody (1:600 or 1:800 in 1% BSA TBS 0.1%TritonX) overnight at 4°C. Next, slides were washed with TBST and incubated with the required fluorescent-dye conjugated secondary antibody (1:1000) for 1 h at RT (Alexafluor, Invitrogen, Thermofisher). Slides were mounted with DAPI Pro-LongDiamond antifade mountant overnight and analyzed the next day. Images were acquired using the Leica SP8 laser scanning confocal microscope and Leica LAS-X software. Analysis of IF images was performed using FIJI, considering the Raw Intensity values.

### Evaluation of PAR levels

Tumor PAR levels were measured using Poly(ADP-Ribose) ELISA Kit (Cell Biolabs, Inc) following the manufacturer’s instructions. Briefly, proteins were extracted in RIPA buffer supplemented with PDD and the PARP1 inhibitor provided by the kit. ELISA plates were coated overnight with the Anti-Poly(ADP-Ribose) coating antibody. All the following antibody incubations were performed for 1 h at RT. Samples were assayed in triplicate (50 µg per well), in parallel with a PAR polymer standard curve, on the pre-coated plate. Next, wells were washed in Wash Buffer x 3 times and incubated with the anti-Poly(ADP-Ribose) Detection Antibody. Following 3 wash steps, samples were incubated with Secondary Antibody-HRP conjugate and washed thoroughly 3 times. Next, 100 µL of substrate solution was added to each well and incubated until visibly developed. After adding the stop solution, absorbance (OD 450 nm) was detected on Envision Excite multilabel microplate reader. Nuclear fractions PAR levels were measured by dot blot (DB). Briefly, 15 µg of SN or CB protein extracts were blotted onto a nitrocellulose membrane gentle vacuum (Bio-rad), and air dried for 15 min. After 1 h of blocking in milk 5% at RT, the membrane was incubated overnight with the anti-PAR antibody at 4°C. HRP-coniugated anti-mouse antibody was incubated 1h at RT. Chemiluminescence signals were acquired via iBright Imaging System (ThermoFisher Scientific).

### RNA-seq and bioinformatic analysis

RNA from tissue samples was extracted with RNeasy Mini Kit (Qiagen), accordingly to the manufacturer’s instructions. Briefly, eight-ten mg of tumor were homogenized with TissueLyser II (Qiagen) in buffer RLT and transferred in a column for RNA isolation and DNA digestion. RNAs’ purity and quality were validated through Nanodrop and TapeStation (Agilent Technologies) by the Wistar Institute Genomics Facility. Next, library preparation was performed using the SENSE mRNA-Seq Library Prep Kit V2 (Lexogen) according to the protocol’s directions and submitted for sequencing (Illumina). Sequenced reads were aligned using RSEM along with Bowtie2 against the human genome (version: hg19) or EBV genome (version: NC_007605.1). Notably, only one sample considered had a percentage of genome alignment below 76%. DESeq2 was used to normalize the reads and obtain the differentially expressed genes between Veh and BMN-673. Genes that passed FDR<5% (p<0.05) threshold were considered significant. Significant genes were then analyzed via Ingenuity Pathway Analysis (IPA). Additionally, enrichment analysis was done using Gene Set Enrichment Analysis (GSEA) on pre-ranked lists generated based on DESeq2 results. STRING clustering analysis on the transcripts codifying proteins was performed with a minimum interaction score confidence of 0.4 (1260, cut-off: FDR<5%, |Z|≥2), filtering the network by functional and physical protein-protein association, co-expression, and co-occurrence in databases. Gene Ontology (GO) enrichment analysis on biological and functional processes was then performed on the clusters. The strength value was considered the Log_10_(Number of Genes _observed_ / Number of Genes _expected_ in a specific network. The dataset is deposited in Gene Expression Omnibus (GEO) under the accession number GSE. Selected human genes up- or down-regulated were validated by qPCR and normalized to 18S values (Figure S2B); oligonucleotides used in this study are available in Table S2.

### ChIP-qPCR

ChIP-qPCR assay was performed according to the Upstate Biotechnology, Inc., protocol as described previously, with minor adjustments (64). Briefly, LCLs were double cross-linked with 1% ethylene glycol bis(succinimidyl succinate) (EGS) for 30 min, followed by 1% formaldehyde for 15 min in constant rotation. DNA was sonicated using the Covaris ME220 Focused-ultrasonicator to generate 200- to 500-bp fragments. DNA-protein complexes were immunoprecipitated with anti-PARP1 C-Term, anti-Myc, or rabbit IgG, eluted, and de-crosslinked overnight. Enriched chromatin was then cleaned up after RNase and Proteinase K treatment. Real-time PCR was performed with a master mix containing 1× Maxima SYBR green (ThermoFisher), 0.25 μM primers, and 1/50 of the ChIP-DNA per well. Quantitative PCRs were carried out in triplicate using the QuantStudio 6 Flex real-time PCR system (Applied Biosystem). Results were analyzed considering the threshold cycles (*C_T_*) by the *ΔΔC_T_* method relative to the Input DNA, and then normalized to the IgG control. The oligonucleotides used in this study are available in SI Appendix, Table S3.

### Quantification of EBV genome copy number

EBV genome copy number was determined using multiplex digital droplet PCR as previously described with minor modifications (65). DNA was extracted from tumors using the DNeasy Blood & Tissue Kit following manufacturer instructions. An equal amount of DNA (100 ng/µl) was digested with HindIII enzyme (New England Biolabs) for 1 h at 37°C. Digested DNA was diluted to a final concentration of 10ng/µl. Samples were prepared by adding 1x of master mix, 1x EBV-LMP1-FAM, and 1x Ribonuclease P protein subunit p30 (RPP30)-VIC probes, to 10 µl of the diluted DNA, per well. Samples were tested in duplicate on a ddPCR plate (Bio-rad). The loaded plate was sealed with aluminum foil (Biorad), and briefly vortexed to homogeneously mix the samples. After centrifugation for 3 min at 1200 rpm, the plate was loaded in the QX200 Droplet Digital PCR (Bio-Rad) automated system to create the droplets. The samples were then transferred to a new plate to run the PCR reaction. Droplet call was executed using QX200 Droplet Reader (Bio-Rad). The EBV copy number was determined considering the concentration of LMP1 positive droplets (copies/volume per well) with respect to the concentration of RRP30 positive droplets divided by 2 (number of RRP30 alleles in the human genome). LCL NHC1 and Akata-BL digested DNA was used as positive and negative control, respectively.

### Statistical analysis

All experiments in this work were conducted at least in duplicate to ensure the reproducibility of results. GraphPad statistical software package was used to identify statistically significant differences between experimental conditions and control samples, using one-way, Two-way ANOVA, Mixed-effects analysis, Mann-Whitney or multiple Student’s t-test as indicated in the figure legends. Outlier analysis was performed using ROUT method.

## Acknowledgments

We also thank James Hayden and Frederick Keeney for scientific support. We are grateful to The Wistar Institute’s Animal, Genomics and Histotechnology core facilities for providing technical support. IT was supported by R01 AI130209, R01 GM124449, and by Core Grant P30 CA010815-53 (PI: Altieri). PML was supported by R01 AI153508, R01 CA259171, R01 DE01733 and by Core Grant P30 CA010815-53 (PI: Altieri).

