## Supplemental data for "PARP1 Inhibition Halts EBV+ Lymphoma Progression by Disrupting the EBNA2/MYC Axis"

Associate Professor, Associate Director for Cancer Research Career Enhancement

Gene Expression & Regulation Program,

Ellen and Ronald Caplan Cancer Center

The Wistar Institute

3601 Spruce Street

Philadelphia, PA 19104, USA

**This PDF file includes:**

Figures S1 to S3 with legends

Tables S1

**Other supporting materials for this manuscript include the following:**

Table S2 with supporting materials and methods

Table S3 Oligonucleotides used in this study.

SI Appendix references

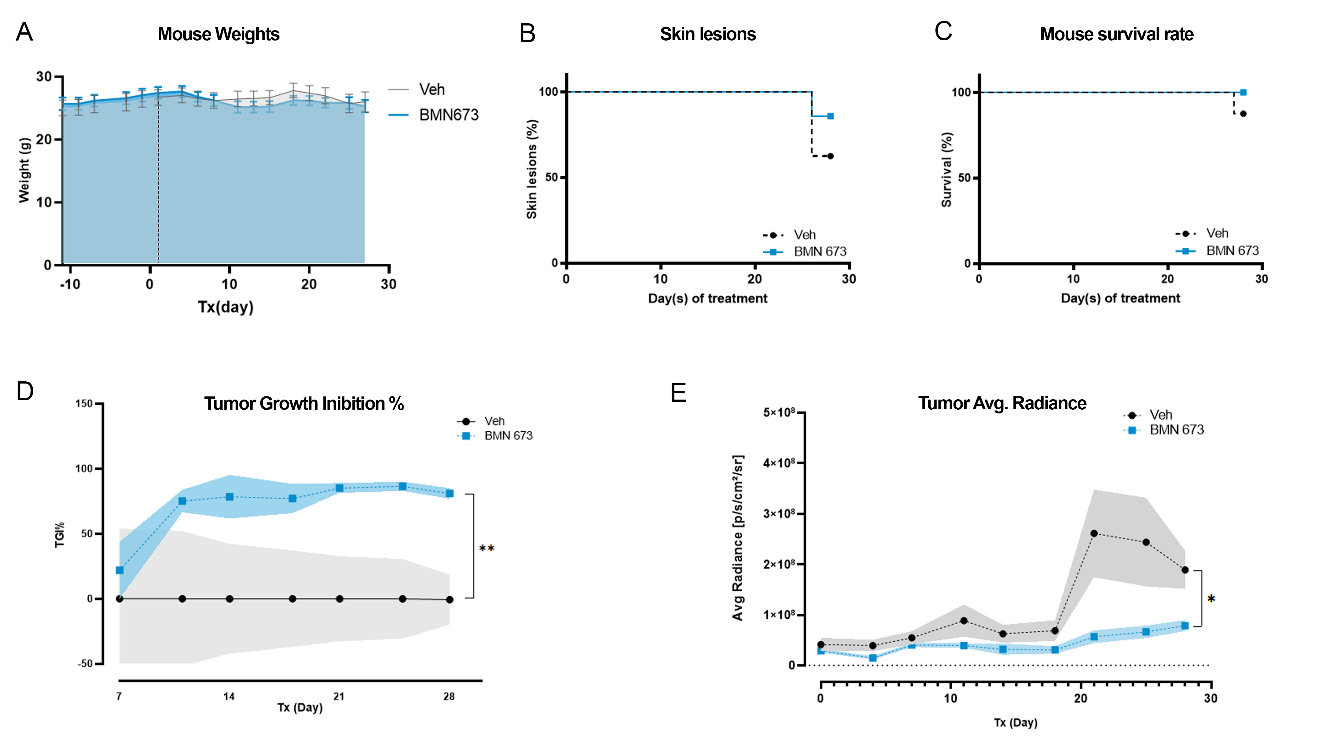
Fig. S1. Mice health and tumor progression during PARP1 inhibitor treatment. (A) Average of mouse weight (g) ± SEM observed throughout the study. (B) Percentage rate of skin lesions (e.g., in the tumor area) during the treatment. (C) Survival rate (%) of the mice within the treatment span. Results show the average (dotted line) ± standard error (SEM; highlighted areas) of the (D) tumor growth inhibition (TGI%, 80.85% ± 4.15, ***p*=0.002), and (E) average radiance ([p/s/cm2/sr]; **p*=0.04) between Veh and BMN 673-treated mice at T_28_ (n=7 Veh; n=8 BMN 673), analyzed by multiple-effects analysis and Sidak *post-hoc* test, or Mann-Whitney test.

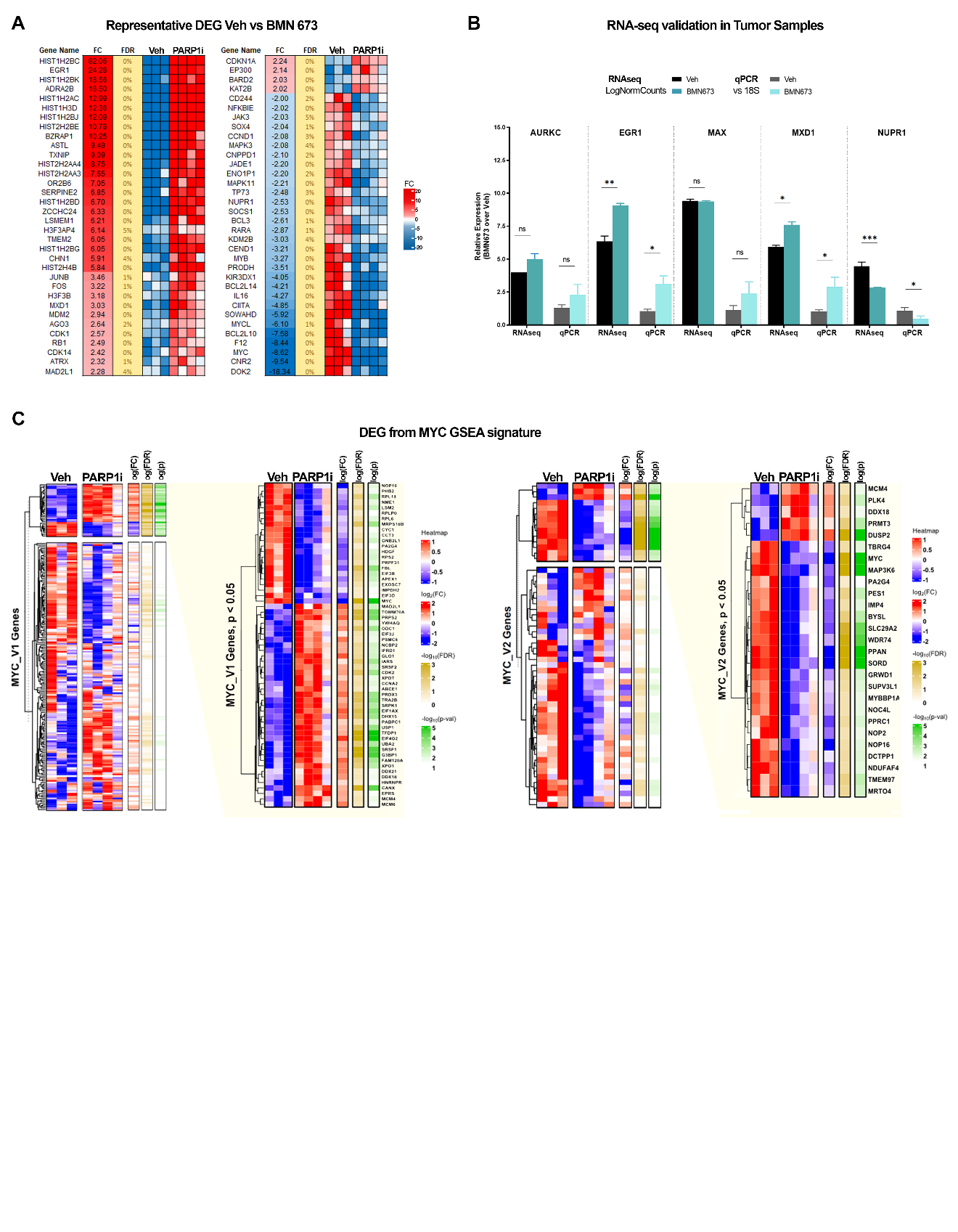

Fig. S2. PARP1 inhibition induces transcriptomic changes. (A) Heatmap of top DEG from Veh and BMN 673 treated samples (PARP1i) showing representative genes that were significantly (FDR≤5%) altered by PARP1 inhibition, ordered by fold-change. (B) RNA-seq validation in tumor extracts. Selected transcriptomic hits are reported and separated by dotted lines. Relative Log normalized expression obtained from DESeq2 for Veh (black) and BMN 673 (teal) are reported on the left bar plots. Gene expression was validated by qPCR and normalized to 18S in both Veh (gray) and BMN673 (cyan)-treated mice, as shown on the right bar plots. Statistical significance was determined by multiple t-test and corrected by two-stage step-up FDR method (EGR1, RNAseq **q=0.002, qPCR *q=0.042; MXD1, RNAseq *q=0.033, qPCR *q=0.033; NUPR1, RNAseq ***q<0.001, qPCR *p=0.034). (C) Heatmaps of genes from MYC GSEA pathway (MYC_V1 and MYC_V2). Significant (p<0.05) genes are highlighted in the yellow magnification.

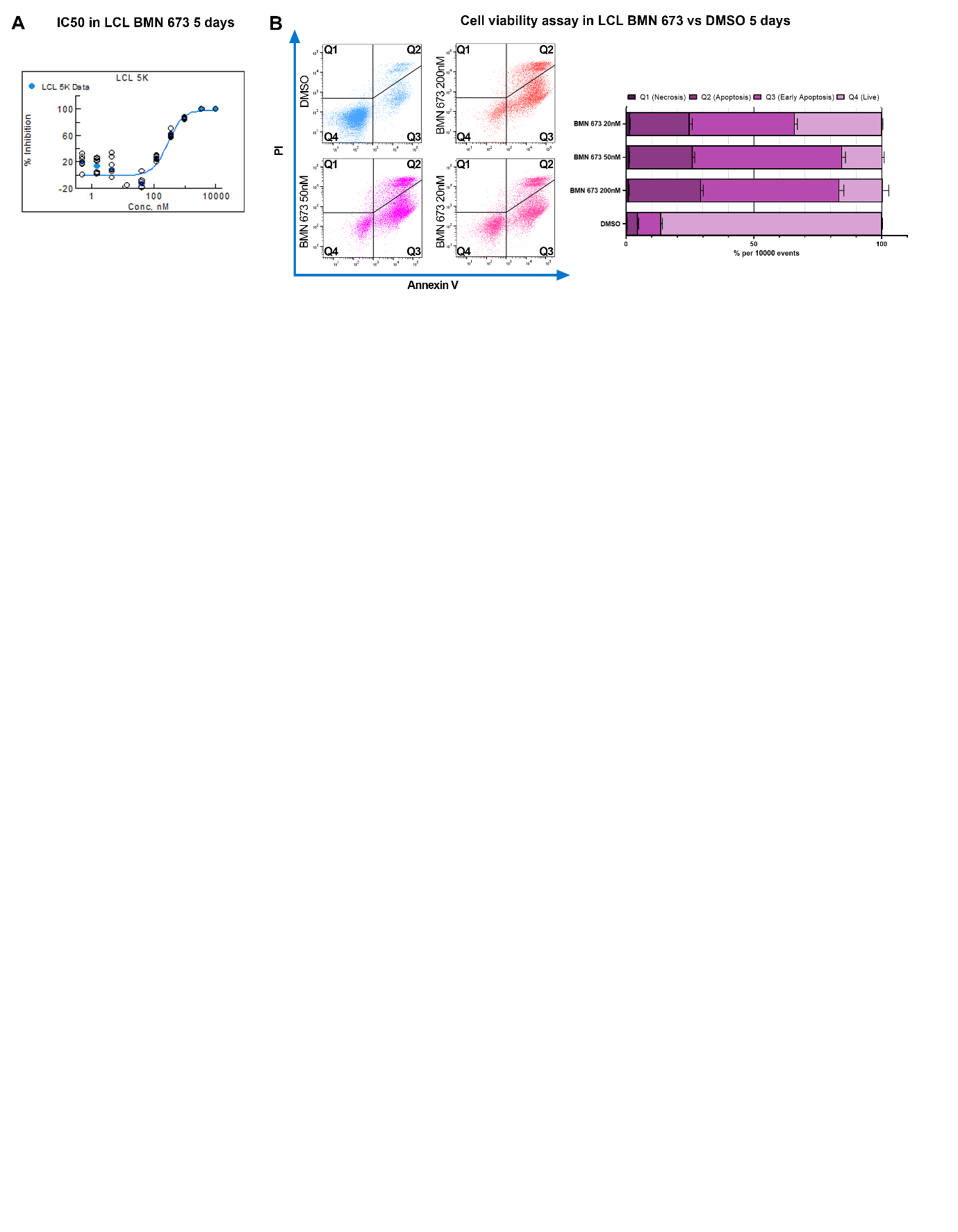

Fig. S3. PARP1 inhibition in LCL. (A) BMN 673 IC50 after 5 days of treatment. Resultant EC50 is ~200 nM. (B) AnnexinV/PI analysis shows the LCL triplicates treated with DMSO (control) or 20 nM, 50nM, or 200nM BMN 673 for 5 days. Ten-thousand events were collected per sample. Quadrants describe PI (Q1, necrosis), AnnexinV/PI (Q2, apoptosis), AnnexinV (Q3, early-apoptosis) positive cells, or negative cells (Q4, live cells).

Table S1. STRING Network GO Biological Function of DEG. Top 20 GO Biological Functions organized by Strenght.

| **GO #term ID** | **GO Biological Processes description** | **observed gene count** | **background gene count** | **Strength** | **FDR** |
| --- | --- | --- | --- | --- | --- |
| GO:0043372 | Positive regulation of CD4-positive, alpha-beta T cell differentiation | 10 | 29 | 0.73 | 0.0147 |
| GO:0042771 | Intrinsic apoptotic signaling pathway in response to DNA damage by p53 class mediator | 10 | 30 | 0.71 | 0.017 |
| GO:1900101 | Regulation of endoplasmic reticulum unfolded protein response | 9 | 28 | 0.7 | 0.0364 |
| GO:0002312 | B cell activation involved in immune response | 12 | 45 | 0.62 | 0.0205 |
| GO:0031295 | T cell costimulation | 14 | 55 | 0.6 | 0.0127 |
| GO:0072332 | Intrinsic apoptotic signaling pathway by p53 class mediator | 13 | 51 | 0.6 | 0.017 |
| GO:0043370 | Regulation of CD4-positive, alpha-beta T cell differentiation | 12 | 47 | 0.6 | 0.0263 |
| GO:0002377 | Immunoglobulin production | 14 | 57 | 0.58 | 0.0147 |
| GO:0045621 | Positive regulation of lymphocyte differentiation | 24 | 102 | 0.56 | 0.00041 |
| GO:0045582 | Positive regulation of T cell differentiation | 21 | 89 | 0.56 | 0.0012 |
| GO:0002762 | Negative regulation of myeloid leukocyte differentiation | 12 | 52 | 0.55 | 0.0455 |
| GO:0008630 | Intrinsic apoptotic signaling pathway in response to DNA damage | 16 | 71 | 0.54 | 0.0133 |
| GO:0045670 | Regulation of osteoclast differentiation | 15 | 67 | 0.54 | 0.0174 |
| GO:0002440 | Production of molecular mediator of immune response | 18 | 83 | 0.53 | 0.0085 |
| GO:1902107 | Positive regulation of leukocyte differentiation | 33 | 156 | 0.52 | 6.98E-05 |
| GO:0045638 | Negative regulation of myeloid cell differentiation | 19 | 97 | 0.48 | 0.0136 |
| GO:0050870 | Positive regulation of T cell activation | 40 | 209 | 0.47 | 6.19E-05 |
| GO:0045580 | Regulation of T cell differentiation | 28 | 147 | 0.47 | 0.0012 |
| GO:1903039 | Positive regulation of leukocyte cell-cell adhesion | 42 | 228 | 0.46 | 6.19E-05 |
| GO:0045619 | Regulation of lymphocyte differentiation | 33 | 178 | 0.46 | 0.00041 |

Table S2. SI materials and methods.

| REAGENT or RESOURCE | SOURCE | IDENTIFIER |
| --- | --- | --- |
| Antibodies | | |
| EA-D  EBV Early Antigen Diffuse Ea-D antibody | Abcam | Cat#ab30541; RRID:AB_732194 |
| EBNA3A | Abcam | Cat#ab16126; RRID: AB_732057 |
| EBNA3B,  Sheep Anti-EBNA 3B Polyclonal Antibody | Abcam | Cat#ab16127; RRID: AB_732058 |
| Anti-Histone H3 antibody | Abcam | Cat#ab1791; RRID: AB_30261 |
| LMP1  EBV Latent Membrane Protein 1 antibody | Abcam | Cat#ab78113; RRID:AB_1566182 |
| P53 | Abcam | Cat#ab1101; RRID: AB_297667 |
| PARP1 | Abcam | Cat# ab191217; RRID: AB_2861274 |
| γH2A.X | Active Motif | Cat#39118; RRID: AB_2793161 |
| PARP1 C-term | Active Motif | Cat#39561; RRID: AB_2793258 |
| PARP1 N-term | Active Motif | Cat#39559; RRID: AB_2793257 |
| Zta (ZEBRA) | Santa Cruz Biotechnology | Cat#sc-53904; RRID: AB_783257 |
| β-Actin (8H10D10) Mouse mAb (HRP Conjugate) | Cell Signaling Technology | Cat#12262; RRID: AB_2566811 |
| MAX | Cell Signaling Technology | Cat#4739; RRID: AB_2281777 |
| MXD1 | Cell Signaling Technology | Cat#4682; RRID: AB_2147734 |
| MYC | Cell Signaling Technology | Cat#13987; RRID: AB_2631168 |
| Goat Anti-Rabbit IgG HRP coniugated | Jackson ImmunoResearch | Cat#111-035-003; RRID:AB_2313567 |
| Rabbit Anti-Mouse IgG HRP coniugated | Jackson ImmunoResearch | Cat#315-035-003; RRID:AB_2340061 |
| Rabbit Anti-Sheep IgG HRP coniugated | Jackson ImmunoResearch | Cat#313-035-003; RRID:AB_2339948 |
| NUMA1 | LSBio | Cat#LS-B11047; RRID:AB_2889921 |
| EBNA2 (clone R3) | MilliporeSigma | Cat#MABE8; RRID:AB_10807963 |
| γH2A.X | MilliporeSigma | Cat#05-636; RRID:AB_309864 |
| PAR | R&D Systems | Cat#4335-MC-100; RRID:AB_2572318 |
| Donkey anti-Rabbit IgG (H+L) Alexa Fluor™ 680 | Invitrogen (ThermoFisher Scientific) | Cat#A-10043 |
| Goat anti-Mouse IgG (H+L) Alexa Fluor™ 594 | Invitrogen (ThermoFisher Scientific) | Cat#A-11005 |
| Chemicals, peptides, and recombinant proteins | | |
| DMSO | MilliporeSigma | D8418 |
| BMN 673 (Talazoparib, LT-673) | Selleck Chemicals | S7048 |
| Etoposide | MilliporeSigma | E1383 |
| PDD | Selleck Chemicals | S8862 |
| Kolliphor-HS 15 | MilliporeSigma | 42966 |
| DMAc | MilliporeSigma | 271012 |
| D-Luciferin, potassium salt | ThermoFisher Scientific | 501532814 |
| Matrigel | Corning | 356231 |
| Proteinase K | MilliporeSigma |  |
| RNase A | ThermoFisher Scientific | EN0531 |
| Formaldehyde solution | MilliporeSigma |  |
| Dynabeads™ Protein A | ThermoFisher Scientific | 10002D |
| Dynabeads™ Protein G | ThermoFisher Scientific | 10004D |
| Bradford assay | Bio-Rad | 5000006 |
| Critical commercial assays | | |
| FITC Annexin V PI Apoptosis Detection Kit | BioLegend | 640914 |
| Comet Assay Kit protocol | Abcam | ab238544 |
| Subcellular Protein Fractionation Kit | Invitrogen (ThermoFisher Scientific) | 78840 |
| EpiQuik™ Total Histone Extraction Kit | Epigentek | OP-0006-100 |
| DNeasy Blood & Tissue Kit | Qiagen | 69504 |
| RNeasy Mini Kit | Qiagen | 74104 |
| Quantseq RNA library prep v2 | Lexogen |  |
| SuperScript™ III First-Strand Synthesis SuperMix | Invitrogen (ThermoFisher Scientific) | 11752250 |
| Pierce™ BCA Protein Assay Kit | Invitrogen (ThermoFisher Scientific) | 23227 |
| Poly (ADP-Ribose) ELISA Kit | Cell Biolabs, Inc | XDN-5114 |
| Experimental models: Cell lines | | |
| LCL NHC1 EBV^+^ (B95.8) |  |  |
| LCL ^mCHERRY/eLUC+^ NHC1 EBV^+^ (B95.8) |  |  |
| Akata Burkitt Lymphoma (EBV^-^) |  |  |
| Experimental models: Organisms/strains | | |
| Mouse: NSG NOD.Cg-Prkdc^scid^ Il2rgtm1Wjl/ScJ | The Wistar Institute Animal Facility |  |
| Software and algorithms | | |
| FIJI | (1) | https://imagej.net/software/fiji/#publication |
| CometAnalyser | (2) | https://sourceforge.net/projects/cometanalyser/ |
| FlowJo v9 | BD Biosciences | https://www.flowjo.com/ |
| Qupath | (3) | https://qupath.github.io/ |
| HALO® IMAGE ANALYSIS PLATFORM | Indica Labs | https://indicalab.com/halo/ |
| R | The Wistar Institute  Bioinformatic core | https://cran.r-project.org/bin/windows/base/ |
| Bowtie2 | (4) | https://bowtie-bio.sourceforge.net/bowtie2/index.shtml |
| DESeq2 | (5) | https://bioconductor.org/packages/release/bioc/html/DESeq2.html |
| ggplot2 | Wickham, 2016 | https://ggplot2.tidyverse.org/ |
| Ingenuity Pathway Analysis | Qiagen | https://digitalinsights.qiagen.com/products-overview/discovery-insights-portfolio/analysis-and-visualization/qiagen-ipa/ |
| GraphPad | Dotmatics | https://www.graphpad.com/ |
| Biorender | Biorender | https://biorender.com/ |
| EndNote X9 | Clarivate^TM^ | https://endnote.com/ |
| Other | | |
| Penicillin-Streptomycin (10,000 U/mL) | ThermoFisher | 15140122 |
| RPMI 1640 | Corning |  |
| radioimmunoprecipitation assay (RIPA) buffer | Millipore | R0278-50ML |
| Laemmli sample buffer | Bio-Rad | 1610747 |

**Table S3. Oligonucleotides used in this study**

| Target | sequence |
| --- | --- |
| CANX_Fw | GCCTCCGCCTCTCTCTTTAC |
| CANX_Rev | CCCCTCTCACCTCTAGCCTC |
| DDX21_Rev | CTTCCCCTCCACGTTCTTTCTT |
| DDX21_Fw | CTGGAGAAGTGGAAGCGAGAC |
| EPRS_Fw | CCAGGGTTAGAACGGCCT |
| EPRS_Rev | CAACCTGCTGTTTTACGACCTG |
| FAM120A_Fw | ACCACATGCTTGGCTACCTG |
| FAM120A_Rev | GCGCCGTTGAAGAAGACGAA |
| PA2G4_Fw | AGCAAACTATCGCTGAGGACC |
| PA2G4_Rev | ACCTTTCCCTATCAGCCTTGA |
| NUPR1_Fw | GACCATGGACACTACACCCA |
| NUPR1_Rev | GAGACTCAGTCAGCGGGAAT |
| EGR1_Fw | GAAAGTTTGCCAGGAGCGAT |
| EGR1_Rev | GGACGGGTAAGAGGTAGCAA |
| AURKC_Fw | AGAAGGAAGGACTGGAGCAC |
| AURKC_Rev | GCTGTGCGCTGTTCATCTAA |
| MXD1_Fw | AAGCTGGGCATTGAGAGGAT |
| MXD1_Rev | CCTGTGAGATAGTCCGTGCT |
| MAX_Fw | CACTCCAAGGAGAGAAGGCA |
| MAX_Rev | GCTGCTCCAGAAGAGCATTC |
